# 3’UTR-directed control of poly(A) tail dynamics and mRNA stability in vertebrate embryos

**DOI:** 10.64898/2026.08.20.746113

**Authors:** Gal Nechooshtan, Mazal Tawil, Talya Razin, Michal Rabani

## Abstract

Post-transcriptional regulation determines mRNA fate through multiple interconnected layers of control, and is particularly important in early embryos. However, how 3’UTR sequences coordinate these different regulatory layers remains poorly characterized. Here, we develop *multi-UTR*, a massively parallel reporter assay that simultaneously tracks poly(A) tail lengths, 3′ terminal nucleotide additions, and mRNA stabilities for thousands of 3’UTR sequences across early zebrafish embryogenesis. We show that embryos use a combination of global and 3’UTR-encoded regulatory programs to progressively remodel mRNA tails. Using reporters with various initial tail lengths, we find that the embryonic cytoplasm rapidly overrides pre-set poly(A) lengths. As development proceeds, 3’UTRs drive tail length diversity and longer tails become progressively associated with increased stability. Strikingly, this association is affected by productive translation: in non-coding reporters, the tail length–stability relationship inverts, such that shorter poly(A) tails are associated with greater stability. Poly(A) tail remodeling is accompanied by two waves of terminal nucleotide additions, early guanylation and later uridylation, that mark distinct regulatory states. Together, our results uncover how the 3′UTR regulatory code operates across multiple layers of regulation to dynamically shape maternal mRNA fate during embryogenesis, and establish *multi-UTR* as a general platform for decoding post-transcriptional regulatory programs.

## Introduction

Cytoplasmic mRNA regulation plays a central role in shaping cellular gene expression programs. Once mRNAs enter the cytoplasm, their fate is tightly controlled by interconnected processes that regulate their stability and translation, with the transcript’s 3’end serving as a central regulatory hub. Poly(A) tail length and tail composition represent two tightly linked regulatory layers. The cellular machinery must both interpret these signals and dynamically remodel them to determine mRNA fate. Modulation of poly(A) tail length is a major driver of mRNA regulation: tail elongation enhances mRNA stability and promotes its translation, whereas shortening has opposite effects ^1–5^. Mechanistically, longer poly(A) tails recruit poly(A) binding proteins more efficiently ^6^, enhancing transcript stability ^7^ and promoting translation through interactions with the translation machinery at the 5’ end ^8^.

In parallel, terminal nucleotide additions provide an additional regulatory layer beyond length. Terminal uridylation marks transcripts for degradation, while the addition of mixed A/G nucleotides shields from rapid deadenylation and subsequent degradation ^9–12^. Viewed together, poly(A) tail length, tail chemistry, and mRNA fate are tightly coupled components of cytoplasmic mRNA regulation.

In early embryos, poly(A) tail dynamics play a particularly dominant role in controlling mRNA stability and translation ^13^. In transcriptionally silent oocytes and embryos, tightly regulated changes to poly(A) length of maternal transcripts control their translation prior to genome activation ^2,9,11,12,14–17^. At those times, long poly(A) tails enhance translation ^2,9,15,16^, while short-tailed mRNAs can remain stable but translationally silent ^18^. At the same time, embryos also couple stability to translation. Inefficiently translated maternal mRNAs enriched in non-optimal codons are targeted for co-translational deadenylation and early decay ^19,20^, driven by CCR4-NOT deadenylases ^21^. This codon-mediated decay works in combination with the mRNA’s 3’ UTR context, such as cis-elements ^19^ and length ^21^. Following genome activation, a massive wave of maternal mRNA degradation clears these transcripts ^22^, and the poly(A) tail regains its canonical role in stability ^16^. The initial translation-dependent decay now acts alongside newly activated zygotic clearance pathways, such as zebrafish miR-430.

mRNA regulation is governed by the interplay of cis-regulatory elements embedded in mRNA sequences, and trans-acting regulators that bind them ^23–28^. Factors such as RNA binding proteins (RBPs) ^26,29–32^ and micro-RNAs (miRNAs) ^33,34^ selectively bind mRNA sequences frequently located within their 3’UTRs, to control transcript fate. For example, binding of zebrafish miR-430 within 3’UTRs induces poly(A) tail shortening and destabilization ^33^, while cytoplasmic polyadenylation element binding proteins mediate poly(A) tail extension to enhance translation and stability ^24^. Systematic studies have mapped global trends and reproducible patterns of 3’ tail changes and mRNA fate ^3,9,16,35–37^. However, genomic analyses struggle to resolve how individual 3’UTR elements shape the fate of the transcripts harboring them within these global trends. Sequence analysis ^38,39^ of endogenous transcripts suffers from a limited sequence diversity, competing effects and combinatorial interactions between co-occurring sequences, and difficulty to isolate the discrete signals that produce complex non-discrete outcomes ^40–42^. Current genomic assays are also still limited in their ability to quantify multiple regulatory processes simultaneously. Massively parallel reporter assays (MPRAs) offer a more controlled setting to mitigate some of these complexities and uncover additional regulatory elements, including those affecting mRNA stability ^43,44^, polyadenylation ^45^ and translation ^45–47^. Injected reporters further circumvent the confounding effect of new transcription. Previously, we developed a reporter system called *UTR-Seq* that successfully decoded sequence-based rules of 3’UTR-mediated mRNA stability in zebrafish embryos ^43^. However, it remains unclear how sequence-based rules within 3’UTRs orchestrate the dynamic, complex and interconnected coordination of poly(A) length remodeling, terminal nucleotide additions, and translational status to ultimately determine mRNA fate.

Here, we develop *multi-UTR*, a dynamic multi-readout MPRA that simultaneously quantifies mRNA abundance, poly(A) tail length, and terminal nucleotide additions for thousands of 3′UTR reporters. Using zebrafish embryos as an in vivo model, we apply *multi-UTR* to reveal how 3’UTR regulatory grammars reprogram 3’ tails to control mRNA fate, and how translational status shapes the functional relationship between 3’ tail states and mRNA stability. Together, our findings establish *multi-UTR* as a powerful platform for dissecting how 3′UTRs encode post-transcriptional regulatory programs during development.

## Results

### *multi-UTR*: a multi-readout MPRA for 3’UTR-directed mRNA regulation in-vivo

To simultaneously dissect several layers of 3′UTR regulation, we developed *multi-UTR*, which reads out poly(A) tail lengths, terminal nucleotide additions, and stability for thousands of mRNA reporters with different 3′UTRs in living embryos.

We used our established 3’UTR reporter system for zebrafish embryos ^43^, in which reporters contain a constant 5′UTR and coding sequence followed by variable 110-bp fragments derived from native zebrafish 3′UTRs. To expand the assay’s readouts, we ligated an adapter directly to the 3’ ends of the mRNA pool after extraction from embryos. We used paired-end sequencing to simultaneously capture reporter identity (3’UTR sequence) and measure mRNA 3’ ends to assess tail changes (**Fig. 1A**). From these reads, we developed a computational pipeline (see **Methods**) to quantify three distinct metrics per reporter (**Fig. 1B**): mRNA abundance, poly(A) tail length, and terminal 3’ end nucleotide additions.

**Figure 1.**
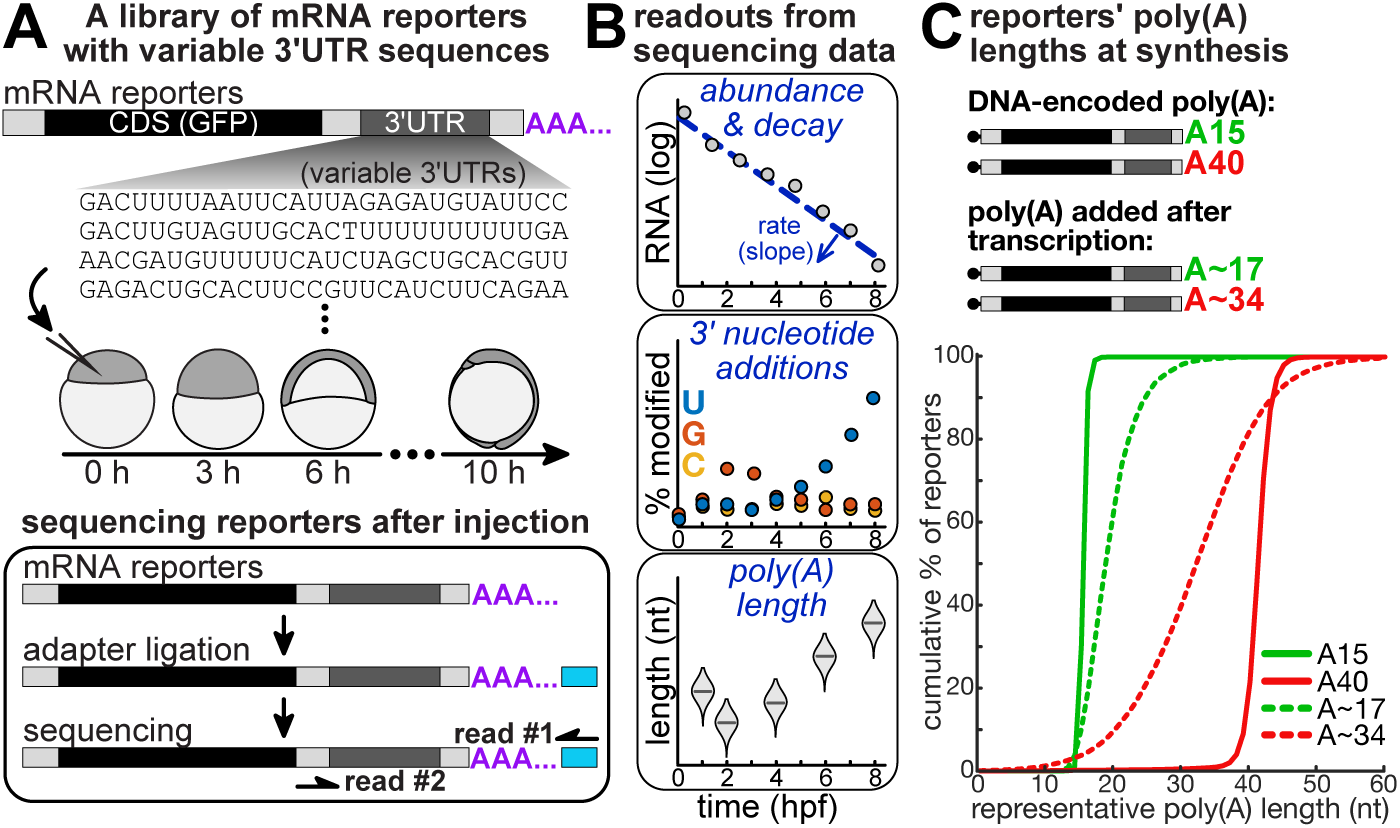
*multi-UTR*: a multi-readout MPRA platform for 3’UTR-based cytoplasmic mRNA regulation *Multi-UTR* is a multi-readout MPRA to assess how 3’UTR sequences regulate cytoplasmic changes in mRNA poly(A) tail length, terminal modification and stability. **(A)** Overview of the *multi-UTR* methodology. mRNA reporters are synthesized in-vitro with a constant backbone, containing a short 5’UTR, a coding sequence (encoding GFP, black), a 3’UTR with variable 110bp sequences (dark gray) and a poly(A) tail (purple; see below). The reporters are injected into zebrafish embryos at the 1-cell stage, and temporal samples are collected along the first 10 hours of development. Spike-ins are added to samples for normalization of both poly(A) lengths and reporter quantities. An adaptor (cyan) is ligated to the 3’ end of extracted RNA and used for library preparation and sequencing of both the reporters’ 3’ end and their 3’UTR sequences (paired-end). **(B)** A computational pipeline quantifies per reporter (bottom to top): representative poly(A) tail lengths, defined by utilizing the 25^th^ percentile of length distribution as a representative in vivo length for each reporter to mitigate right-skewed distributions in high-throughput sequencing of long homopolymeric tracts; fraction with terminal C, G or U nucleotides added at 3’ end; temporal mRNA abundance and resulting degradation rates calculated as the slope of a linear regression line to quantify changes in abundance over time. **(C)** Cumulative distribution (y-axis, fraction) of representative poly(A) tail lengths (x-axis, 25^th^ percentile length measured per reporter) in different reporter libraries. Poly(A) tails are added either as part of in vitro transcription (DNA-encoded; solid lines) or after in vitro transcription, using enzymatic polyadenylation by poly(A) polymerase (dashed lines).

Because native maternal mRNAs exist with diverse, pre-set poly(A) lengths upon fertilization, we designed our assay to determine how initial tails influence mRNA fate. We generated mRNA reporter libraries with distinct initial poly(A) tail lengths (**Fig. 1C**, **Fig S1A**) to reflect different native maternal mRNA populations ^9,16^. These included reporters with precisely encoded and in-vitro transcribed 40nt poly(A) tails (A40), representing highly adenylated maternal mRNAs, and 15nt poly(A) tails (A15), representing the typical poly(A) length of maternal mRNAs at fertilization ^9^. To achieve these precise initial lengths while minimizing the self-templated extensions common in in-vitro transcription, we utilized optimized synthesis strategies, including capture oligos and gapped promoters ^48–50^ (**Fig. S1B**, see **Methods**). These successfully reduced such extensions (**Fig. S1C**), resulting in cleaner ends to both short and long tail libraries. Additionally, to assess transcripts with perfectly clean 3’ ends (**Fig. S1C**), we generated non-adenylated reporters and enzymatically polyadenylated them in vitro to reach mode tail lengths of either ∼34A or ∼17A (**Fig. S1A**).

We injected each of these diverse reporter libraries into one-cell stage zebrafish embryos and collected samples along a developmental timecourse (**Fig. S1D**). Collection started soon after fertilization, proceeded through embryonic genome activation (∼2.5 hours post fertilization, hpf) and continued until the end of gastrulation (∼9hpf). Samples were normalized to internal spike-ins of known poly(A) lengths (20A, 40A) and mRNA quantities, added after extraction.

Thus, *multi-UTR* is designed to interrogate the regulatory logic of maternal 3’UTR sequences in-vivo, combining precisely controlled initial tail lengths and time-resolved multi-readout measurements.

### Global and 3’UTR-directed reprogramming of poly(A) lengths in embryos

To determine how poly(A) tail lengths change after their injection into embryos, we used *multi-UTR* to track a representative tail length per reporter. This measure accounts for known biases in high-throughput sequencing of homopolymeric tracts ^15,37^ (see **Methods**), and confirmed using spike-ins of known poly(A) lengths (**Fig. S2A-B**).

Global poly(A) lengths followed a 3-phase trajectory over time (**Fig. 2A-D**): an initial convergence towards a common length, followed by extension and then renewed trimming, consistent with prior studies ^45^. In the first hour after injection, tail length changes depended on the starting length (**Fig. 2E**, 0-1hpf). Longer initial tails (A40 and A∼34) were trimmed to 23A (**Fig. 2A-B**), while shorter tails (A15) were trimmed only slightly to 13A (**Fig. 2C-D**). Non-adenylated reporters instead had their tails extended over this period (**Fig. S2C-D**, >40% of reporters extended by 1hpf). Between 1 and 4 hpf, tails were globally extended, reaching average lengths of >50A by 4 hpf (**Fig. 2E**, 1-3 hr, 3-4 hr). Trimming resumed after 4 hpf to reach ∼30A by 9hpf (**Fig. 2E**, 4-6 hr, 6-9 hr).

**Figure 2.**
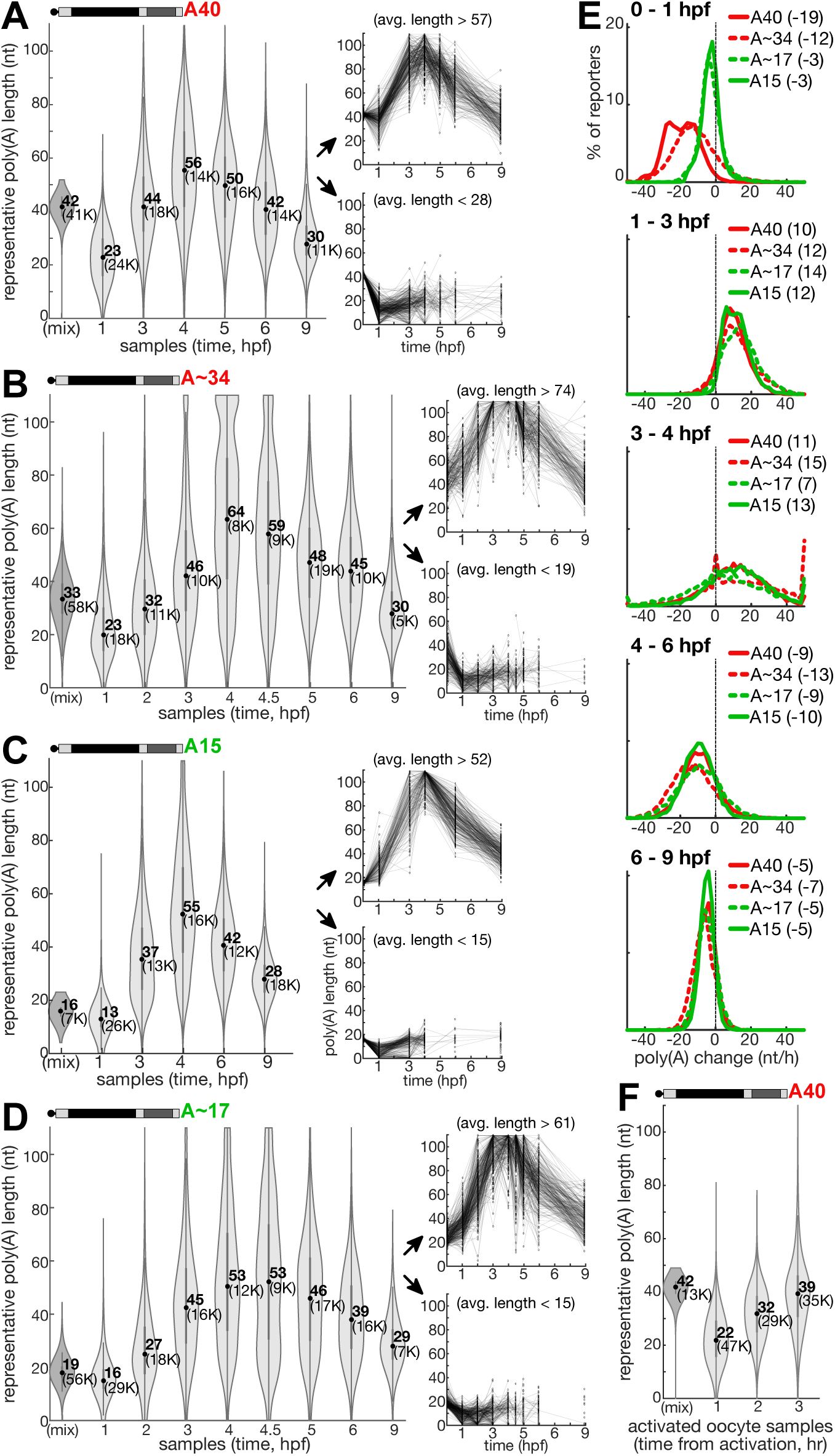
Global and 3’UTR-directed reprogramming of poly(A) lengths in embryos. (A-D) Left: distribution of reporters’ representative poly(A) tail lengths (y-axis, nt, 25^th^ percentile length measured per reporter) in temporal samples (x-axis) of developing zebrafish embryos. The central dot is median; gray central box bounds are 25^th^ and 75^th^ percentiles, upper and lower limits of whiskers are 1.5× interquartile ranges. Values outside of the upper and lower limits are defined as outliers. (mix): in-vitro transcribed RNA library, not injected into embryos. Type of in-vitro transcribed reporter library used is noted on top. Numbers represent mean value, and number of reporters analyzed is noted in brackets. Right: two plots show the representative poly(A) tail lengths (y-axis, 25^th^ percentile length measured per reporter) in temporal samples (x-axis) of a subset of reporters with average poly(A) tail lengths (across all temporal samples) at the top (top plot) or bottom (bottom plot) 2% of the distribution. Panels show results from different reporter libraries: (A) A40, (B) A∼34, (C) A15 and (D) A∼17. **(E)** Distribution (% of reporters, y-axis) of changes in reporters’ representative poly(A) tail lengths (x-axis, nt/h, change is calculated between the two samples noted on top) in developing zebrafish embryos in different reporter libraries. Type of library and mean change is noted in legend. **(F)** Distribution of reporters’ representative poly(A) tail lengths in samples of activated oocytes collected at 1-3 hours after activation. Plots are as defined in (A-D).

Beyond these shared trends, subsets of reporters exhibited noticeably different poly(A) length dynamics (**Fig. 2A-D**, right panels), suggesting that their different 3’UTR sequences directed specific regulation of tail dynamics. Tails of some reporters were extended, while others were trimmed. These divergent behaviors were reproducible across libraries (evident by progressive correlation in tail length, **Fig. S2E**), further indicating they reflect genuine sequence-driven regulation. Thus, embryonic regulation effectively overrides initial poly(A) length differences between libraries, and tunes poly(A) lengths according to 3′UTR context.

To uncover sequences associated with regulation of poly(A) length, we evaluated 4-7nt sequences (k-mers) and identified 449 k-mers with significant differences (FDR-corrected Kolmogorov-Smirnoff p<1%, **Table S1**). By grouping these k-mers based on similarity of both sequence and temporal activity (see **Methods**), we identified a set of 20 clusters, each represented by a single, most significant, k-mer (**Fig. 3**). This analysis uncovered sequence signals, embedded within 3’UTRs, that directed transcript-specific differences in poly(A) length, and acted consistently across reporter libraries. For example, a poly-U motif, matching established cytoplasmic polyadenylation elements ^25,45^, was associated with longer tails (**Fig. S3A**). The miR-430 seed (GCACUU) ^33^ on the other hand, was associated with shorter tails after genome activation (4 hpf compared to 3hpf, **Fig. S3B**). Other elements associated with shorter tails include the AU-rich element (ARE; UAUUUAU, **Fig. S3C**) both before and during genome activation, a C-rich motif (CUCC, **Fig. S3D**) before genome activation, and an A-tract motif (**Fig. S3E**) active shortly after fertilization. These signals broadly matched those identified in previous MPRA studies of poly(A) length ^45,51^ and stability ^43,44^. We further confirmed similar activities (**Fig. S4A-B**) and signals (**Fig. S4C**) in native maternal 3’UTRs^9,16^, consistent with prior gene-specific ^21^ and global studies ^9,16^.

**Figure 3.**
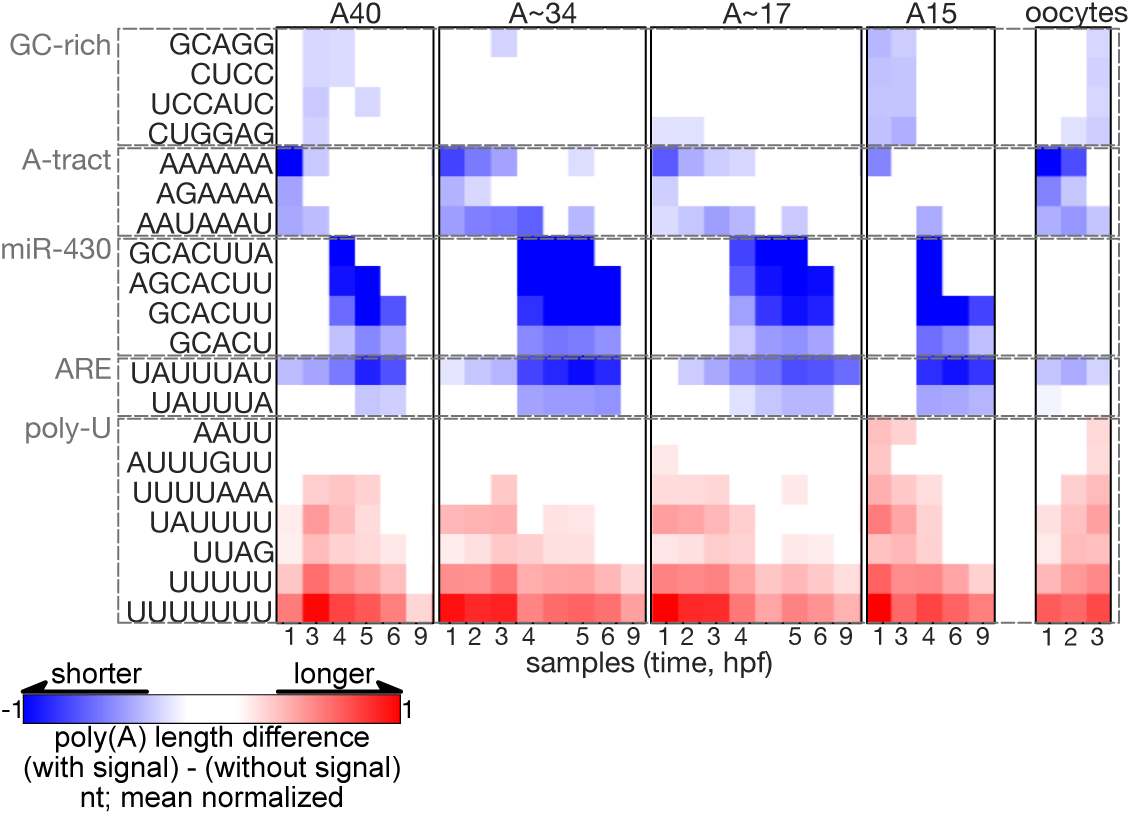
3’UTR sequence signals directing transcript-specific differences in poly. (A) length A set of 20 clusters (rows), each represented by the most significant k-mer, that group sets of enriched k-mers (FDR-corrected Kolmogorov-Smirnoff p<1%) with similar sequence and activity (effect size) across temporal samples and reporter libraries (columns). Colors represent effect size, defined as the mean-normalized difference in average representative poly(A) length in reporters that contain a specific k-mer (signal) and those that do not (blue = negative effect, shorter average poly(A); red = positive effect, longer average poly(A)).

Global trends of early tail shortening and readenylation (**Fig. 2F**) and early signal-specific activities (**Fig. 3**) were also captured in activated oocytes, resulting in correlated poly(A) tail lengths at 3hpf (**Fig. S2F**). Thus, early poly(A) tail reprogramming is driven by maternally inherited activities, that act independently of zygotic factors.

Together, these data show that poly(A) tail length in the embryonic cytoplasm is dynamically changing, and governed by both global activity and 3′UTR-dependent sequence specific regulation.

### 3’UTR sequences direct coupling of poly(A) length to mRNA stability

Next, we asked how 3’UTR sequences shape the relationship between poly(A) tail lengths and stability. We quantified abundance by normalization to known spike-ins amounts added after RNA extraction (**Fig. S5A**), then used those trajectories to estimate degradation rates and validated by comparison to standard *UTR-seq* measurements (**Fig. S5B**).

After injection into embryos, reporters’ abundance decreased over time, with different degradation patterns between reporters. Reporters injected with longer starting poly(A) tails decayed more slowly and with a delayed onset (**Fig. 4A**), as previously observed ^43,51^. However, the degradation rates of individual reporters remained highly correlated between libraries with different starting poly(A) lengths (**Fig. S5C**), indicating that differences in stability between reporters are dictated by 3’UTR context in a similar way despite different starting tails. In all examined libraries, a progressively stronger inverse correlation with stability emerged over time. This coupling began as early as 1 hour after injection (r=-0.45, **Fig. 4B**), peaked by 4 hpf, when poly(A) tails account for up to 48% of the variation in degradation rates (r=-0.69, **Fig. 4B**) and decreased thereafter. Thus, embryos actively reprogram both poly(A) lengths and stabilities according to 3′UTR-encoded regulatory information.

**Figure 4.**
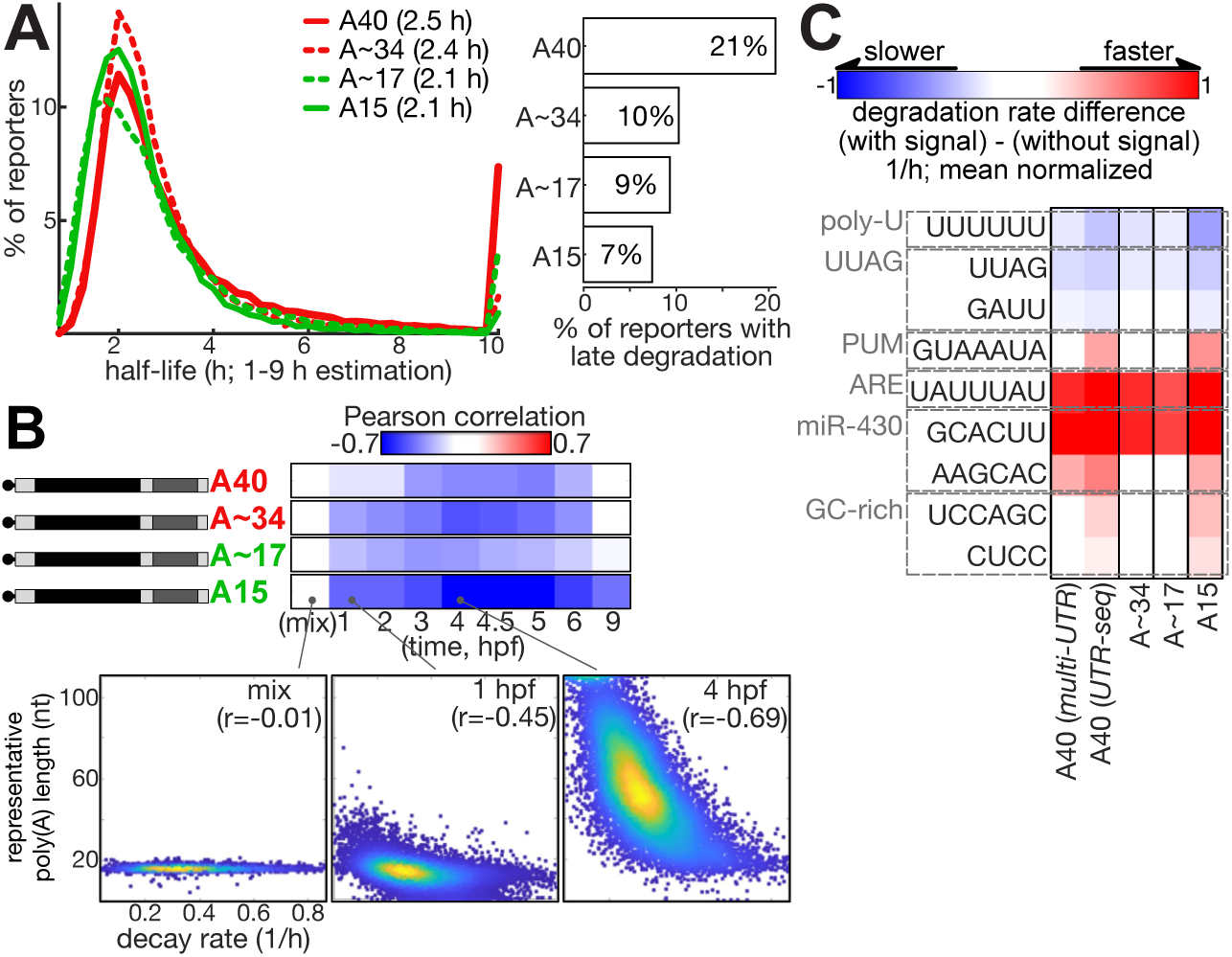
3’UTR sequences direct coupling of poly(A) length to mRNA stability. **(A)** Left: distribution (y-axis, % of reporters) of half-lives (x-axis, h) estimated for reporters in different in-vitro transcribed RNA libraries based on the entire timecourse (1-9 h). Type of library and mean estimated half-life (excluding values larger than upper or smaller than lower bound range; 0.75 – 10 h) is noted in legend. Right: histogram showing % of reporters in each library with late-onset of degradation, identified based on early stability (half-life ≥10 h, estimated on early (1-4 hpf) samples) and late degradation (half-life <10 h, estimated only on late (3-9 hpf) samples). **(B)** Top: Pearson r coefficients (color: red = positive correlation, blue = negative correlation) between estimated degradation rates (x-axis, 1/h) and representative poly(A) tail length of reporters (y-axis, nt, 25^th^ percentile length measured per reporter) in temporal samples (columns) of different in-vitro transcribed reporter libraries (rows). (mix): reporter library before injection to embryos. Bottom: Scatter plots of selected correlations from the A15 library (as indicated by lines). Color represents density (yellow = high density, blue = low density). Pearson r coefficient is noted on top. **(C)** A set of 9 clusters (rows), each represented by the most significant k-mer, that group sets of enriched k-mers (FDR-corrected Kolmogorov-Smirnoff p<1%) with similar sequence and activity (effect size) across reporter libraries (columns). Samples include the 4 *multi-UTR* libraries, and a quantification of the A40 samples also by standard *UTR-seq*. Colors represent effect size, defined as the mean-normalized difference in degradation rates in reporters that contain a specific k-mer (signal) and those that do not (blue = negative effect, slower degradation; red = positive effect, faster degradation).

To identify the specific sequences driving these acquired stabilities, we applied k-mer analysis (see **Methods**) and uncovered 265 significant k-mers (**Table S2**) that clustered into 9 distinct sequence signals (**Fig. 4C**), consistent with previous knowledge ^43,44,51^. Signals remained coherent across libraries, confirming their activity is independent of starting tail length. The regulatory signals affecting mRNA stability (**Fig. 4C**) largely overlapped with those mediating poly(A) length changes (**Fig. 2C**). As expected, stabilizing signals were associated with poly(A) extension, while destabilizing signals were linked to tail trimming, a key step in canonical mRNA degradation pathways. A notable exception was the A-tract motif (**Fig. 2C**), linked to poly(A) trimming but not to any major differences in stability. Instead, this motif was associated with early repression of translation, as evident in genomic ribo-seq data (**Fig. S6**). A Pumilio binding signal (PUM, **Fig. 4C**) was linked to mRNA decay, but not to any major poly(A) length differences, consistent with Pumilio’s known association with some deadenylation-independent destabilization mechanisms ^52,53^.

Together, these results reveal a link between poly(A) tail length and mRNA stability in embryos that builds up over time, and is actively programmed through 3′UTR sequences that shape both properties.

### Two temporally distinct terminal nucleotide additions mark different regulatory regimes

Non-templated terminal nucleotide additions are a second layer of regulation at the 3’ ends of transcripts, beyond tail length. Using *multi-UTR*, we next investigated their in vivo occurrence. We analyzed 3’ end sequences to identify terminal nucleotide additions of U, G, or C bases (≤5 nt, see **Methods**). We focused on enzymatically polyadenylated A∼17 and A∼34 libraries to avoid confounding self-templated extensions during mRNA synthesis, and confirmed the consistency of this analysis on spike-ins (**Fig. S7A**).

Two distinct terminal nucleotide additions emerged in vivo (**Fig. 5A**). First, terminal-U additions increased following genome activation, as was also observed for native maternal mRNAs in zebrafish and other organisms ^9,54^. While these terminal-U additions initially mostly consisted of a single nucleotide, the frequency of longer terminal-U tracts increased over time (**Fig. 5B**). Second, we also identified elevated terminal-G additions prior to genome activation (until 5 hpf). Early guanylation has been shown in human embryos ^11^ but has not been previously observed in zebrafish native maternal mRNAs ^9^. Finally, terminal-C additions were only detected at minimal background levels throughout development. Data from RNA libraries with DNA-encoded poly(A) tails (A40 and A15) showed similar trends (**Fig. S7B**).

**Figure 5.**
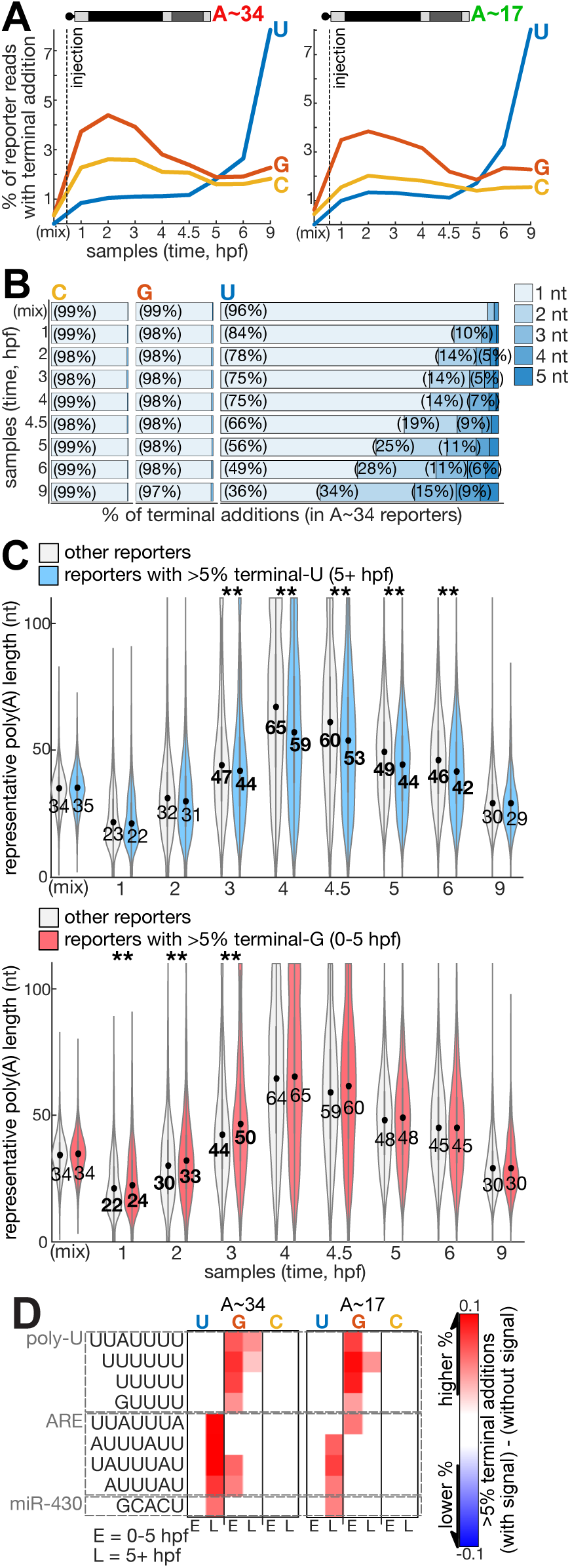
Two temporally distinct terminal nucleotide additions mark different regulatory regimes. **(A)** % of reporter reads with terminal nucleotide additions (y-axis, 3 nucleotides: G = red, C = yellow, U = blue) in temporal samples from zebrafish embryos (x-axis) estimated for reporters in two reporter libraries that were enzymatically polyadenylated with poly(A) polymerase (left: A∼34, right: A∼17). **(B)** Histogram of different terminal nucleotide additions (x-axis, % of terminal nucleotide additions) observed in A∼34 reporters across temporal samples from zebrafish embryos (y-axis). Terminal nucleotides analyzed (left to right): C, G or U. Terminal additions composed of up to 5 consecutive bases of the same base are analyzed (different shades of blue). Numbers on histogram denote % of relevant terminal nucleotide additions. **(C)** Distribution of representative poly(A) tail lengths (y-axis, nt, 25^th^ percentile length measured per reporter) in temporal samples (x-axis) of developing zebrafish embryos, in reporters with >5% terminal nucleotide additions (red = G early (0-5 hpf); blue = U late (5+ hpf)) or reporters with a lower % (gray). The central dot is median; gray central box bounds are 25^th^ and 75^th^ percentiles, upper and lower limits of whiskers are 1.5× interquartile ranges. Values outside of the upper and lower limits are defined as outliers. Numbers represent mean value, bold fonts and ** denote a significant difference by Kolmogorov-Smirnoff test between reporters with >5% terminal nucleotide additions and reporters with a lower %, using a 1% Bonferroni correction. **(D)** A set of 9 clusters (rows), each represented by the most significant k-mer, that group sets of enriched k-mers (FDR-corrected hypergeometric p<1%) with similar sequence and activity (effect size) across two reporter libraries and three terminal bases in each library (columns). Colors represent effect size, defined as the difference in fraction of reporters with >5% terminal nucleotide additions between reporters that contain a specific k-mer (signal) and those that do not (blue: negative effect, depleted in terminal nucleotide additions; red: positive effect, enriched in terminal nucleotide additions).

Crucially, terminal-U and G additions were also coupled to opposing poly(A) tail lengths. Reporters with frequent terminal-U (>5% of reads) possessed shorter poly(A) tails starting at 3 hpf, both on average (**Fig. 5C**) and within individually modified reads (**Fig. S7C**), as previously reported ^37^. Conversely, reporters with frequent terminal-G (>5% of reads) maintained longer average poly(A) tails prior to genome activation (1-3 hpf, **Fig. 5C**). Indeed, mixed A/G tailing has been linked to longer tails in embryos ^11,55^, and has been shown to shield mRNAs from rapid deadenylation ^10^. Despite these longer tails, degradation of reporters with terminal-G was actually faster on average (**Fig. S7D**).

To connect high frequency of terminal nucleotide additions to specific 3’UTR sequence contexts, we used hypergeometric k-mer enrichments (with 1% FDR, see **Methods**). We identified 63 k-mers enriched in modified reporters (**Table S3**) and grouped them into 9 clusters (**Fig. 5D**). Sequence signals overlapped with those previously linked to poly(A) tail changes (**Fig. 2C**) and stability (**Fig. 3C**). For instance, ARE and miR-430 signals, already linked to shorter poly(A) tails and faster decay, were also enriched in terminal-U reporters. On the other hand, U-rich motifs, previously linked to longer poly(A) tails and stability, were enriched in terminal-G reporters.

Together, these data reveal two temporally distinct terminal nucleotide additions marking different regulatory regimes, where uridylation tags shorter tails of unstable transcripts after genome activation, and guanylation is associated with earlier longer tails.

### A massive degradation of non-coding reporters is enhanced by longer poly(A) tails

All reporters characterized so far were translated. To test whether the 3′UTR-directed regulation we observed depends on translation, we generated non-coding reporters lacking AUG triplets in the reporter backbone (see **Methods**). These retain a functional 5’cap that allows ribosomes scanning, but unlikely to support productive translation without an AUG start codon.

Overall, the poly(A) lengths of non-coding reporters (**Fig. 6A**) followed a similar trend to that of their coding counterparts (**Fig. 2A**). However, initial deadenylation of non-coding reporters was enhanced and no final deadenylation was evident (**Fig. 6B**). Poly(A) lengths of non-coding reporters were also linked to similar early 3’UTR signals (**Fig. 6C**) as coding reporters (**Fig. 2C**), but no 3’UTR signals were identified after genome activation, possibly due to technical limitations as a result of their enhanced degradation (see below). Terminal nucleotide additions in non-coding reporters (**Fig S8A**) also follow similar trends as in coding reporters.

**Figure 6.**
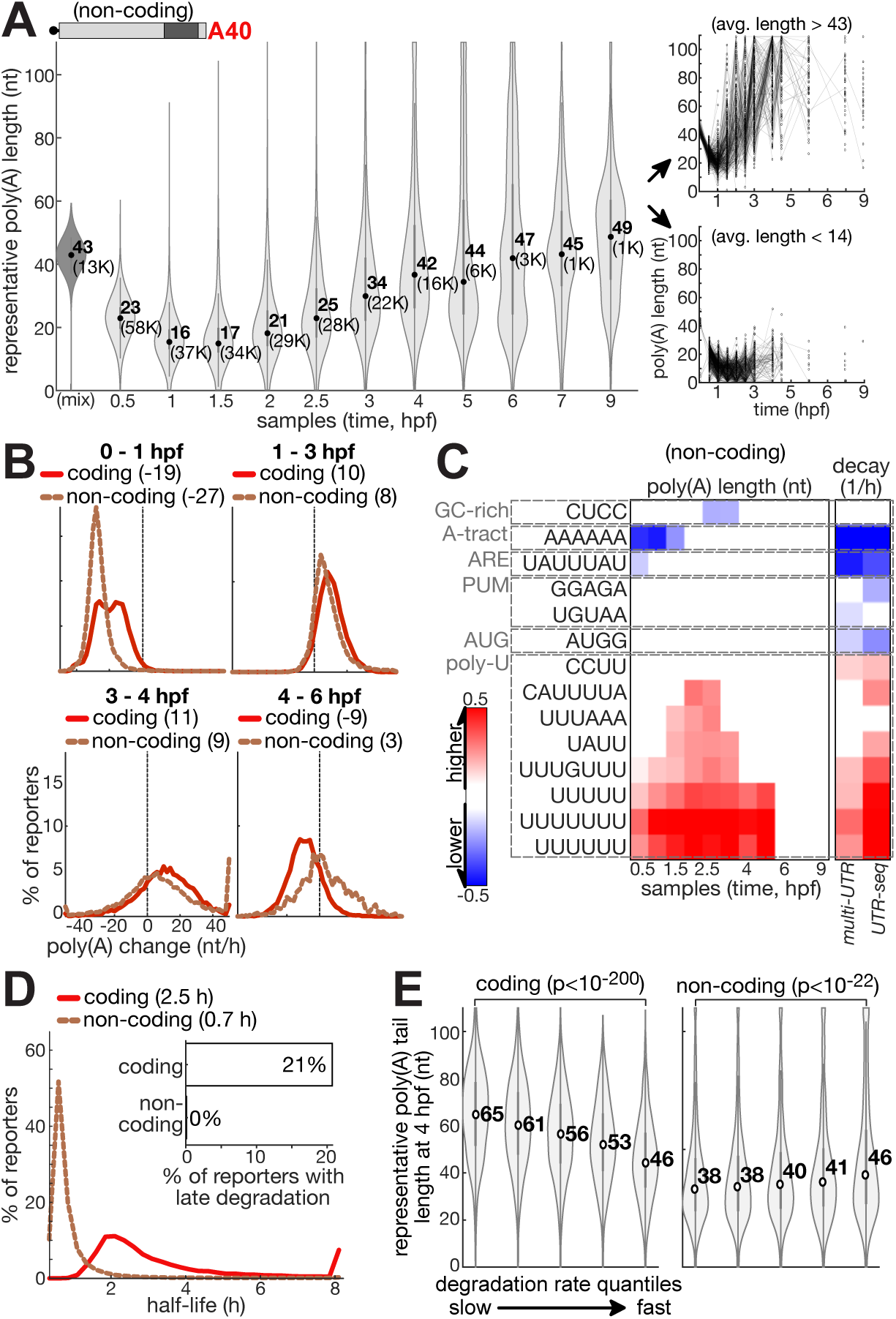
A massive degradation of non-coding reporters is enhanced by longer poly(A) tails. **(A)** Left: distribution of non-coding reporters’ representative poly(A) tail lengths (y-axis, nt, 25^th^ percentile length measured per reporter) in temporal samples (x-axis) of developing zebrafish embryos. The central dot is median; gray central box bounds are 25^th^ and 75^th^ percentiles, upper and lower limits of whiskers are 1.5× interquartile ranges. Values outside of the upper and lower limits are defined as outliers. (mix): in-vitro transcribed RNA library, not injected into embryos. Numbers represent mean value, and number of reporters analyzed is noted in brackets. Right: Two plots showing representative poly(A) tail lengths (y-axis, 25^th^ percentile length measured per reporter) in temporal samples (x-axis) of a subset of reporters with average poly(A) tail lengths (across all samples) at the top (top plot) or bottom (bottom plot) 2% of the distribution. **(B)** Distribution (% of reporters, y-axis) of changes in reporters’ poly(A) tail lengths (x-axis, nt/h, change is calculated between the two samples noted on top) in developing zebrafish embryos in coding and non-coding reporter libraries. Type of library and mean change is noted in legend. **(C)** A set of 14 clusters (rows), each represented by the most significant k-mer, that group sets of enriched k-mers (FDR-corrected Kolmogorov-Smirnoff p<1%) with similar sequence and activity (effect size) across poly(A) lengths or degradation rates by *UTR-seq* and *multi-UTR* protocols (columns), of non-coding reporters. Clustering was applied separately to poly(A) lengths (8 clusters) or degradation rates (7 clusters), and the two final sets of k-mers were merged. Colors represent effect size on representative poly(A) length, defined as the mean-normalized difference in average representative poly(A) length in reporters that contain a specific k-mer (signal) and those that do not; and degradation rates, defined as the mean-normalized difference in degradation rates in reporters that contain a specific k-mer (signal) and those that do not (blue = negative effect; red = positive effect). **(D)** Distribution (y-axis, % of reporters) of half-lives (x-axis, h) estimated for coding and non-coding reporters. Type of library and mean half-life estimated (excluding value outside upper and lower bounds) are noted in legend. Insert: histogram showing % of reporters in each library with late-onset of degradation, identified based on early stability (half-life ≥10 h, estimated on early samples (1-4 hpf)) and late degradation (half-life <10 h, estimated on late samples (4-9 hpf)). **(E)** Distribution of representative poly(A) tail lengths at 4 hpf (y-axis, nt, 25^th^ percentile length measured per reporter) in subsets of reporters divided by their degradation rates (5 equal sized quantiles, x-axis). Left: coding reporters, right: non-coding reporters. The central dot is median; gray central box bounds are 25^th^ and 75^th^ percentiles, upper and lower limits of whiskers are 1.5× interquartile ranges. Values outside upper and lower limits are defined as outliers. P-values are testing the hypothesis of unequal mean between first and last quantiles (Wilcoxon rank-sum test).

Looking at stability, a stark functional divergence emerged: non-coding reporters were highly unstable, exhibiting a 3-fold shorter half-life on average (reduced from 2.7 hr to 0.7 hr, **Fig. 6D**). This massive degradation initiated immediately following their injection, despite a long starting poly(A) tail that stabilized coding reporters. Most strikingly, the coupling of poly(A) tails and stability was inverted in these non-coding transcripts: longer poly(A) tails were linked with faster degradation (**Fig. 6E**). Moreover, sequence analysis showed that early 3’UTR signals inversely affected the stability of non-coding reporters (**Fig. 6C**). Signals associated with longer poly(A) tails, such as poly-U motifs, actually enhanced degradation, while those associated with shorter poly(A) tails, including ARE and A-tract signals, enhanced stability. Interestingly, the presence of an AUG triplet within the 3’UTRs of non-coding reporters was also associated with enhanced stability, likely because it allowed ribosomes to initiate translation and trigger translation-induced stabilization.

Finally, to verify that non-productive ribosome scanning was not responsible for this massive degradation, we synthesized a non-coding reporter library with a non-functional (ApppG) 5’cap, which does not support ribosome recruitment. These non-scanning reporters exhibited the same massive degradation and poly(A) regulation (**Fig. S8B**), confirming that scanning is not the trigger for destruction.

Together, these results uncover a massive degradation program targeting untranslated transcripts. Within this program, the canonical relationship between tail length and stability is inverted, and longer poly(A) tails become associated with greater instability. Thus, translation modulates how the embryo interprets tail length.

### An internal A-tract motif promotes early deadenylation uncoupled from rapid decay

While 3’UTR signals affecting poly(A) length and mRNA stability largely overlapped, an internal A-tract motif within 3’UTRs was a notable exception: associated with poly(A) shortening immediately after fertilization (**Fig. 3**), but not linked to any major differences in stability (**Fig. 4C**). The motif exerted a similar regulatory activity also on native maternal poly(A) lengths (**Fig. S4C**), and was also associated with lower translation of maternal transcripts shortly after fertilization (**Fig. S6**).

High-resolution data (**Fig. 7A**) refined the deadenylation timing of motif-containing reporters to the first 30 minutes after injection, and confirmed that the presence of the motif drives tail shortening significantly beyond the global deadenylation activity observed at those times. Although its activity ended early, poly(A) lengths of motif-containing reporters remained significantly shorter, until ∼4 hpf in most polyadenylated libraries (**Fig. S9A-B**).

**Figure 7.**
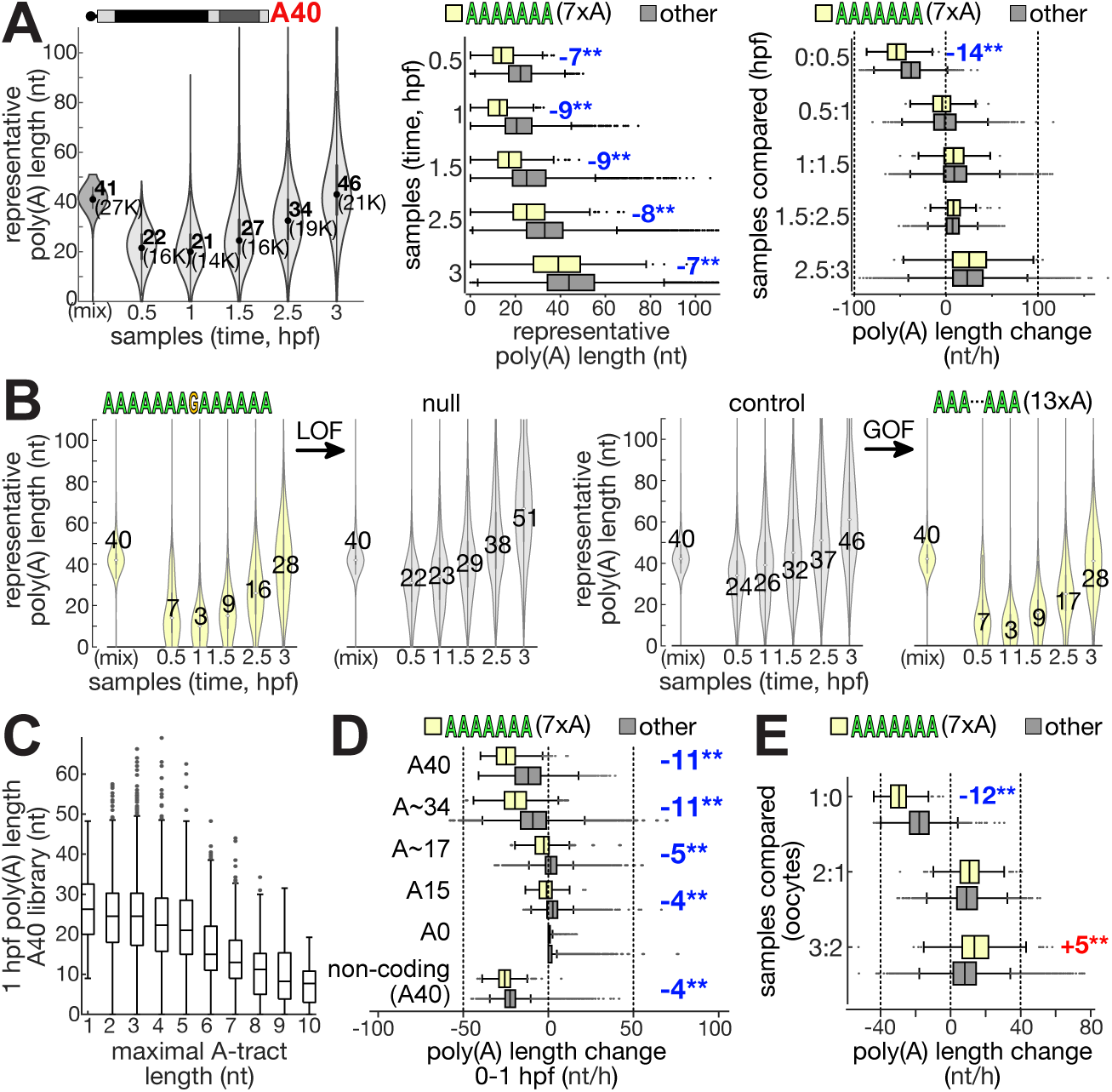
A 3’UTR A-tract motif is associated with early poly(A) shortening. **(A)** Left: distribution of reporters’ representative poly(A) tail lengths (y-axis, nt, 25^th^ percentile length measured per reporter) in temporal samples (x-axis) of developing zebrafish embryos injected with the A40 library. The central dot is median; gray central box bounds are 25^th^ and 75^th^ percentiles, upper and lower limits of whiskers are 1.5× interquartile ranges. Values outside of the upper and lower limits are defined as outliers. (mix): in-vitro transcribed RNA library, not injected into embryos. Numbers represent mean value, and number of reporters analyzed is noted in brackets. Middle: distribution of reporters’ representative poly(A) tail lengths (x-axis, nt, 25^th^ percentile length measured per reporter) with (yellow) or without (gray) an A-tract motif (7 nucleotides or longer) in the same temporal samples (y-axis). Right: distribution of the temporal change in reporters’ representative poly(A) tail lengths (x-axis, nt) with (yellow) or without (gray) an A-tract motif in the same temporal samples (y-axis; time points compared are separated by a colon). Central line represents the median, box edges are 25^th^ and 75^th^ percentiles, whiskers extend to largest/smallest value except outliers; outlier points are plotted individually. Difference in median is noted on plot, and ** denotes a significant difference by a Kolmogorov-Smirnoff test with a 1% Bonferroni multiple hypothesis correction over all tested short sequences and samples (4-7 nt long, 21,760 sequences; p<10^-9^). **(B)** Distribution of representative poly(A) tail lengths (y-axis, nt, 25^th^ percentile length measured per reporter) of validation reporters with or without specific A-tract motifs, as noted on top of each plot, in temporal samples (x-axis) of developing zebrafish embryos. The central dot is median; gray central box bounds are 25^th^ and 75^th^ percentiles, upper and lower limits of whiskers are 1.5× interquartile ranges. Values outside of the upper and lower limits are defined as outliers. Numbers represent mean values. Gray: validation reporters not containing an A-tract motif; Yellow: validation reporters containing an A-tract motif; Left: loss-of-function (LOF) perturbations of an existing A-tract motif. Right: gain-of-function (GOF) mutations creating an A-tract motif in a control reporter that did not originally contain it. **(C)** Representative poly(A) tail lengths (y-axis, nt, 25^th^ percentile length measured per reporter) of A40 reporters with different length of A-tract motif (x-axis) at 1 hpf. Central line represents the median, box edges are 25^th^ and 75^th^ percentiles, whiskers extend to largest/smallest value except outliers; outlier points are plotted individually. **(D)** Distribution of the difference between reporters’ representative poly(A) tail lengths at the first hour after injection (x-axis, nt) with (yellow) or without (gray) an A-tract motif, for different reporter libraries (y-axis). Difference in median between the other and A-tract group is noted on plot. Plots as in (A). **(E)** Distribution of the temporal change in reporters’ representative poly(A) tail lengths (x-axis, nt) in temporal samples of activated oocytes (y-axis; colon separates time point compared) with (yellow) or without (gray) an A-tract motif. Plots as in (A). Difference in median between the other and A-tract group is noted on plot (blue: A-tract is lower, red: A-tract is higher).

Loss-of-function perturbations and gain-of-function insertions (**Fig. 7B**) of the A-tract motif confirmed it is necessary and sufficient to enhance deadenylation, with only a minimal destabilizing effect (**Fig. S9C-D**). Deadenylation activity scaled with the length of the A-tract motif itself, acting more potently when the motif was longer (**Fig. 7C**, **Fig. S9D**). Similar to global early deadenylation, A-tract directed deadenylation was also stronger in reporters injected with longer initial poly(A) (**Fig. 7D**), and also affected non-coding reporters, albeit with a smaller effect.

A-tract motif associated deadenylation was also captured in oocytes (**Fig. 7E**, **Fig. 3**), and was limited to the first hour after activation. Between 2-3 hpf, the A-tract motif exhibited an inverted regulatory behavior in oocytes, associating instead with enhanced readenylation (**Fig. 7E**), as also observed for other poly(A) length regulatory motifs in embryos (**Fig. S3**).

These results reveal a specialized A-tract motif within 3′UTRs that promotes rapid, targeted deadenylation immediately following fertilization or oocyte activation, uncoupled from degradation.

## Discussion

In this work we develop and apply *multi-UTR*, a multiplexed reporter assay for simultaneously tracking poly(A) tail length, terminal nucleotide additions, and mRNA stability. Using this platform, we show that global trends, specific 3’UTR-driven regulation, and translational status all modulate the functional relationship between transcripts’ 3’ tails and stability during the zebrafish maternal-to-zygotic transition.

### Dynamic poly(A) length changes reflect global trends and 3’UTR-specific regulation

Starting poly(A) tail lengths of reporters changed dynamically after injection, uncovering a global pattern of deadenylation and readenylation (**Fig. 2**). Beyond the global trends, initially homogeneous reporter populations progressively diverged into characteristic poly(A) tail length distributions dictated by their 3′UTR sequences. These findings demonstrate that the embryonic machinery progressively aligns the mRNA physical state with its sequence-based regulatory identity. More broadly, they show that the cytoplasm is a dynamic regulatory environment capable of executing complex and dynamic post-transcriptional programs even in the absence of transcription. This fundamental regulatory logic may extend to other transcriptionally restricted contexts, such as mitosis and neuronal synapses. Applying *multi-UTR* in this context would reveal how 3′UTRs encode post-transcriptional regulatory programs across diverse biological systems, and whether a similar regulatory logic is used.

Poly(A) length changes initiated with a rapid deadenylation that strongly depended on starting tail length. Within 1 hour from fertilization, long starting tails (A40, A∼34) shortened to ∼22nt average length, while shorter starting tails (A15, A∼17) changed much less, reaching ∼13nt average tail length. Notably, endogenous maternal mRNAs have a median poly(A) tail length of 13.5 nt at fertilization ^9^, coinciding with the footprint protected by a single poly(A) binding protein molecule ^56^ from deadenylation by PARN, a central deadenyase in oocyte maturation^57^. Thus, initial deadenylation of longer tailed reporters, which we also identify in activated oocytes, may reflect a similar PARN activity inherited from the oocyte.

Initial deadenylation, was not only affected by starting tail length, but was also regulated by the 3’UTR context. Interestingly, within the first hour of development, both long terminal poly(A) tails and long internal A-tract motifs triggered rapid deadenylation (**Fig. 7**). We speculate that both features act as competitive landing pads for a shared trans-acting regulator. One such factor could be poly(A) binding protein ^7,58^, which subsequently directs deadenylation machinery to the 3′ terminus. Because binding avidity scales with length, longer tracts recruit this regulator more efficiently. However, these two binding sites lead to different results. At the terminal poly(A) tail, the A-tract is both a binding site and a substrate of regulation, resulting in a homeostatic loop that converges to a pre-defined poly(A) length. This regulation can help maintain short poly(A) tails characteristic of maternal mRNAs, and is possibly inherited from the oocyte. On the other hand, the internal A-tract is topologically separated from the 3′ terminus, making it a permanent regulatory fixture shifting transcripts into a short-tailed, stable “storage” state that tunes translation (or other cytoplasmic behaviors) without immediately triggering destruction. A similar A-tract motif activity in non-coding transcripts allows them to mitigate the massive degradation program that targets untranslated RNAs. Thus, a conditional interpretation of poly(A) tails provides a mechanistic framework for the unique behavior of the internal A-tract motif.

### Translation inverts the coupling between mRNA stability and poly(A) tail length

We identify coupling between poly(A) tail length and mRNA stability that builds up in embryos over time (**Fig. 4**). Transcript stability became progressively correlated with tail length starting before genome activation, peaking shortly after zygotic genome activation and subsiding later. These results agree well with current knowledge on the embryonic mode of action. Central factors in the maternal mRNA clearance machinery, such as TUT4/7 and decapping enzymes, are inactive after fertilization ^9,12,18^. Thus, even mRNAs with very short tails are initially protected from degradation. Toward zygotic genome activation, these factors become active, and degrade mRNAs if their tail is not protected. However, protective poly(A) binding proteins are inherited in limiting concentrations from oocytes ^2,6^, allowing transcripts with longer tails to compete better for their protection, and creating an association between tail length and stability. As poly(A) binding proteins start accumulating after zygotic genome activation, their concentration eventually reaches saturating levels and allow transcripts with all but the shortest tails to be protected.

However, tail-stability coupling is not universal. Our study uncovered a surprising translation-dependent inversion of this link (**Fig. 6**). While longer poly(A) tails are associated with stability in coding mRNAs, as the dogma holds ^3^, non-coding transcripts undergo rapid degradation that is faster when poly(A) tails are longer. Indeed, non-coding reporters degraded 3-fold faster than their coding counterparts, consistent with the established instability of untranslated mRNAs ^59^, and the transient nature of non-coding RNAs ^60^. Our data rules out non-productive ribosome scanning as a trigger for destabilization. Our results also differ than what has been reported for inefficiently translated mRNAs ^21^, where blocking of translation with morpholino oligonucleotides triggered mRNA stabilization, possibly due to inhibiting inefficiently translating ribosomes. Our noncoding reporters are probably not subject to such decay, as they are not translated. Despite their fast decay, 3′UTR programs in non-coding reporters still remodeled poly(A) tails similarly to their coding counterparts, but the functional outcome was inverted: in the absence of translation, longer poly(A) tails were associated with faster degradation. Thus, the machinery responsible for early poly(A) remodeling operates similarly with or without translation, while translation promotes mRNA survival. Several mechanisms may explain this inversion. Longer tails better recruit poly(A) binding proteins, which can stimulate mRNA decay ^7,58^, in addition to their more canonical protective activity. In the absence of translation, the former may dominate. Alternatively, in the absence of translation, short-tailed transcripts may benefit from oocyte stabilization mechanisms aimed to protect dormant deadenylated messages in oocytes and early embryos by specialized protein shielding, physical compartmentalization ^61^, or uncoupling of decay pathways ^62^. For example, zebrafish eIF4E1b preferentially associates with the cap of short-tailed maternal mRNAs, promoting their protective localization to cytoplasmic granules ^61^. Future studies will clarify the mechanisms involved in this inversion, and determine the relevance of the non-coding regulatory logic to native embryonic bona fide lncRNAs ^63,64^.

### Terminal nucleotide additions mark distinct regulatory regimes

Alongside poly(A) length adjustments, we identified two temporally distinct phases of terminal nucleotide additions (**Fig. 5**). Consistent with previous studies ^9,12^, terminal uridylation was enhanced after genome activation, and coupled with shorter poly(A) tails. We also identified early terminal guanylation prior to genome activation. This terminal-G was associated with longer poly(A) tails, aligning with recent observations that G/A mixed tailing can shield mRNAs from rapid deadenylation ^10,11^. While specific 3’UTR signals were linked to higher frequency of these terminal additions, those signals largely overlapped the signals affecting poly(A) tail length. Thus, it is unclear if signals directed terminal additions or if additions represent intermediate states within tail length pathways. Current knowledge positions terminal-U as such intermediate state, tagging transcripts for destruction and directed by shorter tails. Terminal-G, on the other hand, appears to be recruited to specific transcripts by 3’UTR sequences. For example, the association of guanylation with poly-U elements (**Fig. 5D**) could reflect a direct mechanism, as these elements are likely bound by CPEB in zebrafish embryos^45^, and CPEB was shown to recruit TENT4B ^65,66^ that executes mixed tailing ^10^.

### The *multi-UTR* methodology

Post-transcriptional regulation is a multi-faceted process. By coupling multiple regulatory readouts, *multi-UTR* provides an integrated view of cytoplasmic mRNA metabolism, that minimizes confounding differences such as cellular context and developmental stage, and improves sensitivity. Biologically, introducing exogenous reporters directly into the cytoplasm bypasses the need to control for confounding cellular processes, such as uneven or variable transcription, but may also change regulation compared to native transcripts. Furthermore, while the platform provides high temporal resolution, it measures averaged behavior across cell populations and does not directly resolve cell-type–specific regulation within the embryo. Although high-throughput sequencing of long homopolymeric sequences is prone to known biases ^15,37^, computational post-processing corrects for them in the generally short embryonic poly(A) tails measured here, allowing precise poly(A) length estimates. Adapting the methodology for long-read sequencing technologies ^17,67,68^ can further enhance measurement accuracy, and even wider applicability.

Together, our findings define a dynamic 3′UTR-encoded regulatory code that orchestrates multiple layers of post-transcriptional regulation, revealing how regulatory sequences program maternal mRNA fate during embryogenesis and providing a framework for studying RNA regulation in other biological contexts.

## Methods

### Zebrafish

All protocols and procedures involving zebrafish were approved by the Hebrew University Ethics Committee (IACUC; Protocols #NS-15859 and #NS-23-17226). Embryos were grown and staged according to standard procedures ^69^. Zebrafish embryos from wild-type AB/TL strains were used for all experiments.

### Design and cloning of 3’UTR reporter plasmid libraries

The *UTR-seq* 3’UTR reporter library ^43^ was used in this study. The library contains a set of 90,000 oligonucleotides that were designed to cover 3’UTR sequences of zebrafish embryonic genes, and cloned into a GFP construct driven by an SP6 promoter, with a short 5’UTR (48nt) and a constant 3’UTR sequence (66nt) flanking the cloning site of the 90,000 sequences. The non-coding reporter library was constructed by mutating all AUG triplets within the reporters’ backbone in all three reading frames, resulting in a total of 12 point-mutations. The resulting 780nt DNA fragment was in-vitro synthesized (IDT gBlocks). Construction was performed by Gibson assembly (NEB #E2611L) of the DNA fragment with an empty *UTR-seq* vector, and validated by sequencing. The *UTR-seq* 3’UTR sequence library was cloned into the 3’UTR of the resulting non-coding vector after linearization with EcoRV via Gibson assembly (NEB #E2611L) with the original *UTR-seq* cloning site adaptors. Bacterial transformations were performend by electroporation into Endura electrocompetent cells (Lucigen) according to the manufacturer’s recommendations. Colonies grown overnight from transformation were harvested and plasmids extracted with EZNA Plasmid Mini (Omega).

### Cloning of poly-A-motif test reporters

Individual wild type and mutated reporters were generated in two PCR steps. In the first step, the 110nt variable fragment, flanked by upstream and downstream 20 nt *UTR-seq* universal adapters, was generated by overlap extension PCR. This reaction included forward and reverse primers (**Table S4**) carrying the *UTR-seq* universal adapter sequences (GGAGATCTGAGTTCAAGGAT and GACTCACTATAGTTCTAGAT, respectively) at their 5’ end and overlaps to a third primer, located at the middle of the fragment, at their 3’ ends. A subsequent PCR reaction amplified the first PCR product with primers annealing to the universal adapters and adding 15nt flanking sequences identical to sequences flanking the universal adapters on the *UTR-seq* plasmid. The products of this reaction were cloned into an empty *UTR-seq* vector linearized with EcoRV using Gibson assembly (NEB #E2611L). Reaction products were used to transform NEB 5-alpha Competent *E. coli* cells (New England Biolabs). The sequence of resulting plasmids was verified by Sanger sequencing.

### DNA standards

DNA standards with 20A, 40A and 65A were amplified from plasmids harboring spike-in reporters with relevant poly(A) tail lengths. Amplification was performed with an upstream indexed library amplification primer annealing to the *UTR-seq* C1 adapter located upstream of the cloned 110nt variable region (**Table S4**), and a downstream primer (#274, **Table S4**) annealing to vector sequence immediately downstream of the poly(A) tail. These primers add the required Illumina adapters. DNA standard with 15A was constructed in two steps. First, a spike-in reporter was amplified with an upstream indexed library amplification primer and a downstream primer (#275, **Table S4**) annealing to vector sequence immediately upstream of the poly(A) sequence and to 15 A residues of the poly(A) sequence. This primer adds a 15nt handle for annealing of a second primer (#274, **Table S4**) immediately downstream of the 15nt poly(A). Following cleanup, this product was used as template in a PCR reaction with the same indexed upstream library amplification primer and downstream primer (#274, **Table S4**). PCR products were sequenced on Illumina NextSeq and NovaSeq platforms.

### Templates for RNA synthesis

Templates for *in vitro* transcription of mRNAs with 40A tails were amplified by PCR from the original *UTR-seq* reporter plasmid library ^43^, the non-coding reporter library, or plasmids harboring sequences of individual reporters. All PCR products were treated with DpnI after amplification to remove plasmid templates. Reporter libraries with 40A tails and 40A tail spike-ins (used as length standard) were transcribed under high salt conditions, to minimize self-templated extensions of RNA ^50^. PCR was performed with Q5U Hot Start DNA polymerase (New England Biolabs), using a forward primer harboring an SP6 RNA polymerase promoter with deoxyuridine at position –4 relative to the transcription start site (#222, **Table S4**) and reverse primer (#300, **Table S4**). PCR products were purified using a DNA Clean & Concentrator kit (Zymo Research) and/or AMPure XP beads (Beckman Coulter), then treated with Thermolabile USER II Enzyme (New England Biolabs). Reactions were inactivated by incubating 10 min at 65°. Templates for transcription of mRNA libraries with 0A tails were amplified with primers #216 and #223 (**Table S4**). Templates for transcription of mRNA libraries with 15A tails were amplified with primers #216 and #273 (**Table S4**). Template for transcription of a spike-in with 20A tail was prepared by linearizing the corresponding plasmid with SapI, which cleaves the plasmid at the end of the poly(A) sequence. The reaction was purified with a DNA Clean & Concentrator kit.

### RNA synthesis

All *in vitro* transcription reactions were performed with the HiScribe SP6 kit (New England Biolabs). Heat inactivated USER reactions were used as template without purification, in reactions containing SP6 Reaction Buffer, 5 mM each of ATP, UTP and CTP, 1 mM of GTP, 4 mM of cap analog (New England Biolabs cat. no. S1411S) or ApppG analog (New England Biolabs cat. no. S1406S), 1.6 U/µl of RiboLock RNase inhibitor (Thermo Scientific), template (0.025-0.035 µM), 0.2-0.3 M of NaCl and 0.1 volume of SP6 RNA Polymerase Mix. Reactions were incubated 1 h at 37° and cleaned up with an RNA Clean & Concentrator kit (Zymo Research) using the manufacturer’s protocol for RNAs ≥200 nt. RNA was treated with TURBO DNase (Invitrogen) and cleaned up as after transcription. For reporters in the poly-A-motif test set, a pool of PCR products from individual reporters was used as template. Reporter libraries with 0A and 15A tails were transcribed in the presence of a 3’-end capture oligo ^49^. Reactions contained SP6 Reaction Buffer, 5 mM each of ATP, UTP and CTP, 1 mM of GTP, 4 mM of cap analog, 25-30 nM of PCR-generated template and 0.1 volume of SP6 RNA Polymerase Mix. Transcription reactions of 15A RNA included 3.5 µM of oligo #272 (**Table S4**). Transcription reactions of 0A RNA included 7 µM of oligo #350 (**Table S4**). RNA was cleaned up and treated with DNase as described above.

Spike-in RNA with 20A tail (to be used as tail length standard) was transcribed and treated with DNase using the HiScribe SP6 kit according to the manufacturer’s manual without cap analog, then cleaned up with an RNA Clean & Concentrator kit. For in-vitro transcription of other spike-in RNAs with a 40A tail, suitable plasmids were linearized with SapI (NEB # R0569L) and resulting products were transcribed and treated with DNase using the HiScribe SP6 kit according to the manufacturer’s manual, and cleaned up with an RNA Clean & Concentrator kit. The length and integrity of all RNAs were tested by analyzing on denaturing polyacrylamide gels.

### In-vitro polyadenylated RNA synthesis

To prepare libraries of reporter RNAs polyadenylated with poly(A) polymerase, 0A RNA libraries prepared as described above were polyadenylated in 20 µl reactions containing 15.5 pmol of RNA, 5 units *E. coli* poly(A) polymerase (New England Biolabs cat. no. M0276S), poly(A) polymerase buffer and ATP (0.035 mM for ∼20A library, 0.15 mM for ∼32A library). Reactions were incubated at 37° for 5 min, then transferred to ice and inactivated by adding 5 µl of 50 mM EDTA. Polyadenylated RNAs were cleaned up with an RNA Clean & Concentrator kit, using the manufacturer’s protocol for RNAs ≥200 nt. To deplete shorter-tailed RNAs, RNA polyadenylated in the presence of 0.035 mM of ATP was poly(A)-selected with the NEBNext poly(A) mRNA magnetic isolation module (New England Biolabs cat. no. E7490), followed by cleanup on a Zymo RNA Clean & Concentrator kit and concentration on a SpeedVac.

### Embryo microinjection and sample collection

Zebrafish embryos were injected at the 1-cell stage with 1 nl of a solution containing 0.05% phenol red, 0.1 M KCl and 100 ng/µl of mRNA library. Following injection, embryos were collected into fresh culture medium (5mM NaCl, 0.17mM KCl, 0.33mM CaCl_2_, 0.33mM MgSo_4_, 0.25mM HEPES, 0. 0001% Methylene blue), grown and staged according to standard procedures. 20-25 embryos were collected at selected time points after visual inspection as detailed for each individual experiment (**Fig. S1C**).

### Oocyte collection and microinjection

Mature oocytes were collected by squeezing the belly of adult female zebrafish under anesthesia with tricaine solution (0.15 mg/ml). Oocytes were incubated in culture medium for 5 min. to induce maturation. Oocytes were injected after maturation and remained incubated in culture medium until collection. A total of 5-10 oocytes were randomly collected per sample.

### RNA isolation

Collected embryos were resuspended in 400 µl of fresh culture medium and 100 µl of 5 mg/ml pronase (Roche) were added. After 5 min incubation at room temperature, culture medium was removed without exposing embryos to air, and 500 µl of TRI Reagent (Sigma-Aldrich) was added. Remaining injection mix was also resuspended in 500 µl of TRI Reagent. Embryos were disrupted by pipetting multiple times, incubated 5 min at room temperature and transferred to – 80°. RNA isolation proceeded according to the manufacturer’s protocol.

### Added spike-ins

For *Multi-UTR* of 0A and 40A timecourse samples, 5.55 pg of a mix of equal parts of 40A spike-in (**Table S4**) synthesized at high salt and 20A spike-in was added per 1000 ng of total RNA. For all other library preparations, spike-ins included 40A spike-in synthesized at high salt, 20A spike-in, and 3 additional 40A spike-ins with distinct 3’UTR sequences (**Table S4**), mixed in a ratio of 8:8:4:2:1, respectively, was prepared. For *UTR-seq* of 0A and 40A timecourse samples, 1.6 pg of the mix was added per 1000 ng of total RNA. For *Multi-UTR* of samples from 15A, A∼17 and A∼34 timecourses, 8 pg of the mix was added per 1000 ng of total RNA. For non-coding reporters timecourse samples, a mix of spike-ins (in a ratio of 16:8:4:2:1) was added to the embryos in TRI Reagent (0.2 pg per embryo). Before adapter ligation, RNA was further supplemented with 5.55 pg of a mix of equal parts of 40A spike-in and 20A spike-in per 1000 ng of total RNA.

### *Multi-UTR* library preparation

RNA was ligated to a pre-adenylated, 3’-blocked DNA adapter, in 10 µl reactions containing 600 ng of total RNA, T4 RNA ligase buffer, 20 units of RiboLock RNase inhibitor, 25% PEG 8000, 5 µM of adapter and 200 U of T4 RNA ligase 2, truncated KQ (New England Biolabs). RNA isolated from remaining injection mixes and source RNA from *in vitro* transcription reactions were also ligated to adapter. Ligation reactions were incubated overnight at 16°, then cleaned up with an RNA Clean & Concentrator kit. Adapter-ligated RNA was reverse transcribed with a primer (#239, **Table S4**) annealing to the ligated adapter. Primer annealing was performed in 14.5 µl reactions containing adapter-ligated RNA, 2 µl of 10 µM primer and 1 µl of 10 mM dNTPs. Reactions were heated to 65° for 5 min, followed by cooling to 50°. Then, 4 µl of 5× reverse transcription buffer, 0.5 µl (20 U) RiboLock RNase inhibitor and 1 µl (200 U) of Maxima H Minus reverse transcriptase (Thermo Scientific) were added. Reactions were incubated 30 min at 50°, 5 min at 85° and cooled to 4°. cDNA was used as template in PCR reactions with primers annealing to the *UTR-seq* universal adapter upstream of the 110 nt variable fragment and to the reverse transcription primer, and carrying barcoded Illumina adapter sequences. Amplification was performed with LongAmp Hot Start Taq (New England Biolabs), using a combined annealing and synthesis step at 57°. Reactions were purified with AMPure beads and sequenced on Illumina NextSeq or NovaSeq platforms.

### *UTR-seq* library preparation

RNA was reverse transcribed with a primer (*UTR-seq* RT primer, **Table S4**) annealing to the *UTR-seq* universal adapter downstream of the 110 nt cloned variable fragment. Primer annealing was performed in 6.5 µl reactions containing 1367-1500 ng of total RNA and 1 µl of 10 µM primer. Reactions were heated to 65° for 5 min, followed by cooling to 4°. Then, 1 µl of 10 mM dNTPs, 2 µl of 5× reverse transcription buffer and 0.5 µl (100 U) of Maxima H Minus reverse transcriptase were added. Reactions were incubated 50 min at 50°, 5 min at 85° and cooled to 4°. RNA isolated from remaining injection mixes and source RNA from *in vitro* transcription reactions were also reverse transcribed. cDNA was used as template in PCR reactions with primers annealing to the *UTR-seq* universal adapter upstream of the 110 nt variable fragment and to the reverse transcription primer, and carrying barcoded Illumina adapter sequences. Amplification was performed with LongAmp Hot Start Taq, with primer annealing at 52° and extension at 65°. Reactions were purified with AMPure beads and sequenced on Illumina NextSeq or NovaSeq platforms.

### *Multi-UTR* sequencing data processing

#### Sequencing reads were processed by the following procedure

<u>Read #1 processing (3’end of the molecule)</u>. Information on poly(A) tail length and terminal nucleotide additions was recorded using the following procedure. (1) Trimming the RT adapter 3’end sequence (17nt) using cutadapt ^70^. Untrimmed reads were discarded. (2) Trimming the RT adapter 5’end sequence (8nt) using cutadapt. Untrimmed reads were discarded. Sequence preceding the trimmed adapter was retained as UMI (12bp). (3) Searching the maximal poly-T stretch that starts within the first 25nt of the trimmed read, with upto 5% non-T bases. Stretches of 10 or more bases that had sequencing quality < 15 were considered non-T bases. This is the raw “poly(A) tail length”. Sequences preceding the poly(A) tail on the read is the raw “terminal nucleotide addition”. (4) Searching for the *UTR-Seq* C2 constant sequence (20nt) within the trimmed read using cutadapt. If not found, and raw poly(A) tail length < 8, the read is considered “illegal” and removed from the analysis.

<u>Read #2 processing (5’end of the molecule)</u>. Information on reporter identity was recorded using the following procedure. (1) Trimming the *UTR-Seq* C1 constant sequence (20nt) using cutadapt. Untrimmed reads were discarded. (2) Trimming the *UTR-Seq* C2 constant sequence (20nt) using cutadapt. (3) If the sequence left after trimming < 5nt, reporter is considered “empty” (no insert). (4) Otherwise, align the trimmed sequence to a reference set of all 90,000 synthetic oligonucleotide sequences using Bowtie2 ^71^ (‘end-to-end’ and ‘very-sensitive’ parameters, edit score of 20 or less, quality score of 10 or more). Information of matching reporter was recorded.

<u>Merged data</u>. Only fragments for which both read #1 and read #2 were successfully analyzed were retained for further poly(A) and terminal nucleotide additions analysis. Each fragment contains information on poly(A) tail length, terminal nucleotide additions, UMI and reporter identity. Reads with the same combination of UMI and reporter identity were considered duplicated, and filtered. The most frequent poly(A) tail length and matching terminal nucleotide additions were retained.

<u>mRNA abundance</u>. Information on reporter identity from read #2 processing was combined with UMI information from read #1 processing, for reads that passed two steps of RT adapter trimming. For each reporter identity, number of unique UMIs identified per sample was collected into count matrices used for downstream processing.

### *Multi-UTR* reporters’ poly(A) tail lengths estimation

Merged data from each sample was analyzed to excludes any reads with any terminal nucleotide additions. Any poly(A) lengths above 120nt were capped at 120nt. All reads for each unique reporter were grouped, and representative poly(A) tail lengths for each reporter were estimated using a quantile-based approach. As intragenic poly(A) lengths were shown to approximate a right-skewed lognormal distribution ^15^, and possibly further enhanced by technical noise in high-throughput sequencing of long homopolymeric tracts ^37^, we defined the representative poly(A) tail length of each reporter as the 25th percentile (1st quartile) of all valid reads mapped to a given reporter. This conservative metric anchors the measurement to the dense, lower end of the distribution, granting robust protection against right-tail technical noise while preserving high sensitivity targeted tail shortening. Only reporters with at least 10 valid reads were retained for further analysis.

### *Multi-UTR* reporters’ mRNA decay kinetics prediction

Only reporters with an average count of 10 UMIs per sample across the entire timecourse were analyzed. Raw UMI reporter counts per samples were adjusted relative to the total UMI depth of the sample to calculate transcripts per million (TPM) values. Only expression values based on at least 8 UMIs were analyzed. TPM values across all temporal samples within an experiment were normalized relative to internal spike-in RNA controls, to reach an absolute scaling over time. A liner regression (matlab *lsqlin* function) of temporal log-transformed normalized expression values was applied to calculate degradation rates. Degradation rate was defined as the negative value of the regression slope, using lower and upper bounds to ensure the slope remains negative (representing decay rather than synthesis) and falls within biologically plausible limits. Lower bound is 0.02 RNA/hr (half-life equivalent of ∼35 hr) and upper bound is 5 RNA/hr (half-life equivalent of ∼10 min.). The models evaluated the fit using the sum of squared residuals to calculate the Mean Squared Error (MSE) and the R-squared value. For calculating “early” degradation rates, we fitted only samples collected by 4hpf. For calculating a “late” degradation rate, we fitted only samples collected at 3hpf and later. Final degradation rates lower than 0.07 (half-life equivalent of ∼10 hr) were floored to 0.07.

### *Multi-UTR* reporters’ terminal nucleotide additions frequencies estimation

Merged data from each sample was filtered to excludes any reads with a poly(A) tail shorter than 4nt or a terminal nucleotide addition longer than 5nt, both of which likely reflect sequencing, base-calling or tail read processing errors rather than genuine tail structure. Retained reads were classified by the composition of their terminal addition. The large majority (70-74%) consisted of a homopolymeric addition of 1-5 repeats of a single nucleotide type (U, G, or C), and were classified accordingly. A smaller fraction (17-21%) contained a single non-A nucleotide type mixed with residual A bases. Because our poly(A) tail-calling procedure (Read #1 processing) tolerates up to 5% non-A bases within the called tail itself, such reads cannot be reliably distinguished from tail-calling noise and were excluded. The remaining ∼9% contained a combination of at least two of U, G, or C and were excluded due to their lower frequency and ambiguous composition. Reporters covered by fewer than 10 reads were also excluded. For each reporter, we calculated the fraction of reads carrying each terminal nucleotide addition (U, G or C) out of its total read count, and capped any frequency below 0.5% at 0.5%.

### *UTR-seq* data analysis and decay kinetics prediction

Analysis of *UTR*-seq sequencing reads was performed as previously described ^43^. Analysis of the UMI counts obtained from these samples was performed using the same procedure as described above for the *multi-UTR* decay kinetics prediction.

### Sequence k-mer enrichment analysis

<u>Continuous properties</u>. We associated a short sequence (k-mer) with a regulatory effect when reporters that contain this k-mer in their 3’UTR had a significantly different distribution (one-sided Kolmogorov-Smirnov test, 1% FDR) of a specific continuous property (poly(A) length, degradation rate etc.) than reporters without this sequence. We assigned an effect size to each k-mer by calculating the standardized mean difference defined as 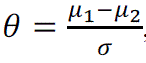, where *μ*_1_ is the mean value of the tested property within the first population, *μ*_2_ is the mean within the second population and *σ* is the standard deviation (based on both populations). We tested all short sequences (k-mers) between 4-7 nucleotides long.

<u>Discrete properties</u>. We associated a short sequence (k-mer) with a regulatory effect when reporters that contain this k-mer in their 3’UTR were enriched for the occurrence of the regulatory effect (terminal nucleotide additions frequency >5%) relative to reporters without this sequence (one-sided Hypergeometric test, 1% FDR). We assigned an effect size to each k-mer by calculating the difference in the frequency (a 0-1 scale) of a regulatory effect within the two populations. We tested all short sequences (k-mers) between 4-7 nucleotides long.

<u>Poly(A) length associated k-mers</u>. We aggregated significant k-mers across all temporal samples within four reporter libraries (A40, A∼34, A∼17 and A15), and retain only k-mers with significant p-value in more than a single sample, and an average significant p-value below 10^-5^. Retained k-mers were clustered based on a greedy network graph analysis. Graph nodes represent k-mers, and edges are drawn between “similar” k-mers. We define the functional similarity between k-mers as the Euclidean distance between the effect-size vectors (across all samples) of the two k-mers. We define the sequence similarity between k-mers as the edit distance between the two k-mers. The edit distance is 0 when one k-mer is contained within the other k-mer. Based on these two metrics, we define two k-mers as “similar” if functional distance is at the lower 50% of all distances calculated, and the edit distance is at most 3. The greedy network graph analysis selects at each iteration the most significant k-mer remained, and clusters it with all other connected k-mers in the graph. This k-mer is the selected representative of the cluster.

<u>Terminal nucleotide additions associated k-mers</u>. We aggregated significant k-mers across two reporter libraries (A∼34 and A∼17), 3 terminal nucleotides (U, G and C) and two temporal windows (0-5 hr, and 5+ hr). We applied the same procedure as with poly(A) lengths to cluster significant k-mers.

<u>Degradation rates associated k-mers</u>. We aggregated significant k-mers across four reporter libraries (A40, A∼34, A∼17 and A15), using degradation rates estimated within 3 temporal windows (all samples, early <=4hpf samples or late 4+hpf samples). We applied the same procedure as with poly(A) lengths to cluster significant k-mers, using the Pearson correlation distance instead of Euclidean distance. For the A40 library we used degradation rates estimated by both *multi-UTR* and *UTR-seq* analysis.

<u>Non-coding reporters’ associated k-mers</u>. We applied the poly(A) length associated k-mers procedure to poly(A) lengths across temporal samples, and the degradation rate associated k-mers procedure to degradation rates estimated from *multi-UTR* and *UTR-seq* counts. We merged the two sets of k-mers by removing redundant k-mers that occurred in both sets.

### Genomic ribo-seq data processing

Zebrafish ribo-seq data and translation-efficiency calculations were taken from ^63^, and sequence motif analysis was performed using QUANTA ^51^.

### Genomic poly(A) tail length data processing

Zebrafish poly(A) tail length data was taken from ^9,16^, and sequence motif analysis was performed using QUANTA ^51^.

## Data and code availability

The code used to generate figures and analyses is openly available at https://github.com/rabanilab/multi-utr.

## Funding

This research was supported by the European Research Council Horizon 2020 (grant 852451 to MR) and by the Israel National Science Foundation (grant 1176/21 to MR).

## Author contribution statement

MR and GN conceived and designed the study. GN developed the *multi-UTR* assay, and performed experiments with help from MR, MT and TR. TR performed oocyte extraction. MR performed data analysis and modeling. All authors read and approved the manuscript.

## Acknowledgements

We thank YD, EM, SS and MMRP for critical reading of the manuscript. We thank Alex Schier for supporting early aspects of this project and for helpful discussions. We thank the Rabani and Nachmani labs for stimulating discussion and feedback throughout our work on the project. We thank Yoel Poloniecki for help with zebrafish maintenance.

---

**Figure S1.**
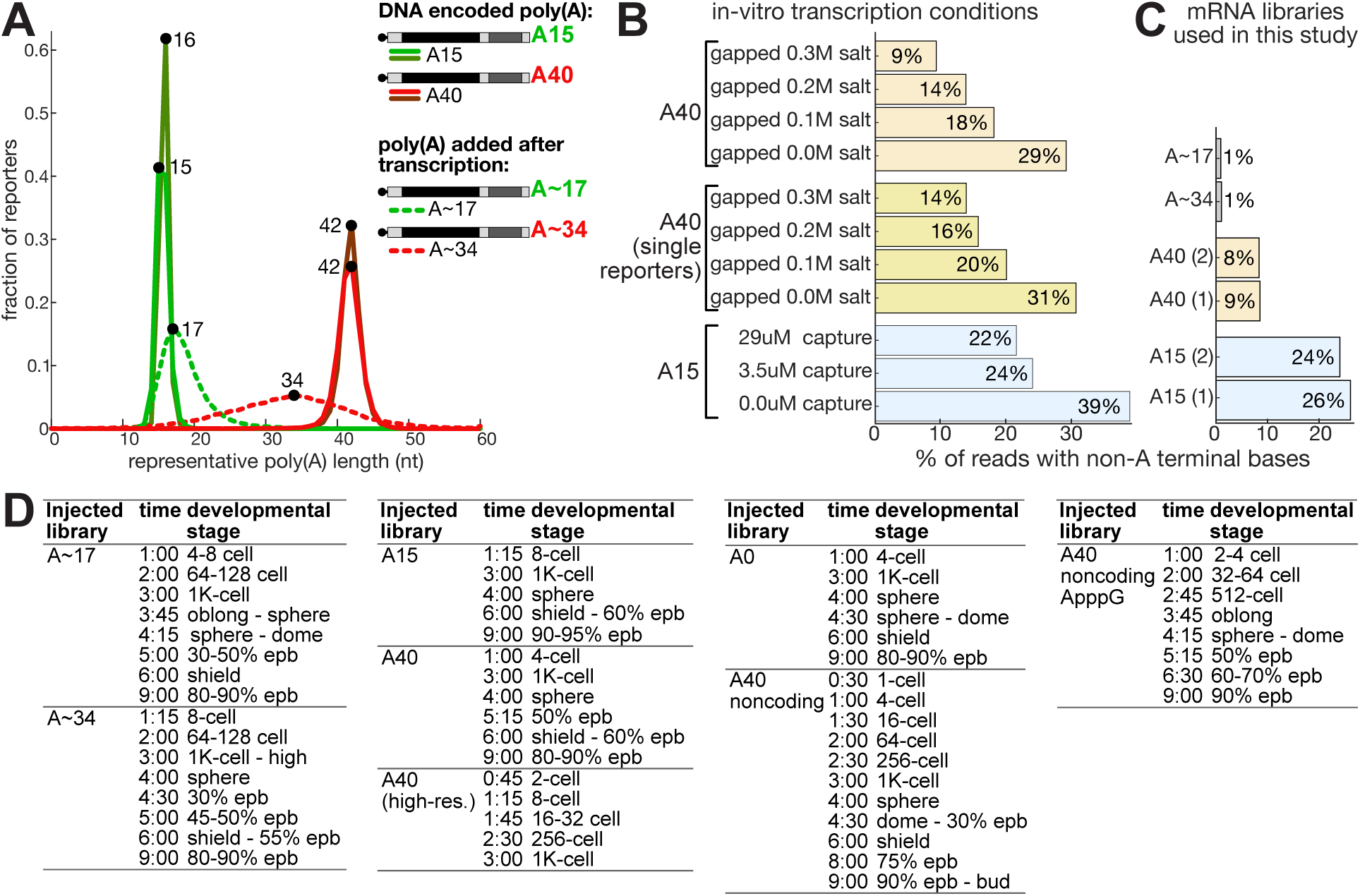
mRNA reporter libraries and collection timelines. **(A)** Distribution (y-axis, fraction) of representative poly(A) tail lengths (x-axis, 25^th^ percentile length measured per reporter) in different reporter libraries. The most frequent value (mode) in each distribution is maked by a black dot and the value is indicated. Two replicates for DNA encoded poly(A) reporters are plotted by different shades of red (A40) or green (A15). **(B-C)** Fraction of reporters with non-A bases identified after the expected poly(A) tail end of transcripts in in-vitro synthesized reporter libraries. **(B)** Fraction of non-A additions using different in-vitro transcription conditions. A40 libraries (top) or single reporters (middle) transcribed with a gapped promoter and increasing salt concentrations (yellow/orange). A15 libraries transcribed with increasing concentrations of capture oligo (bottom, blue). Increased capture oligo or salt concentrations reduced the fraction of reads with non-A additions. **(C)** Fraction of non-A additions in in-vitro transcribed RNA libraries used in this study. A40 libraries (orange) and A15 libraries (blue) are measured across 2 replicates each. **(D)** List of zebrafish embryo samples collected and developmental stages at collection (epb = epiboly). Times are derived from developmental stage at collection.

**Figure S2.**
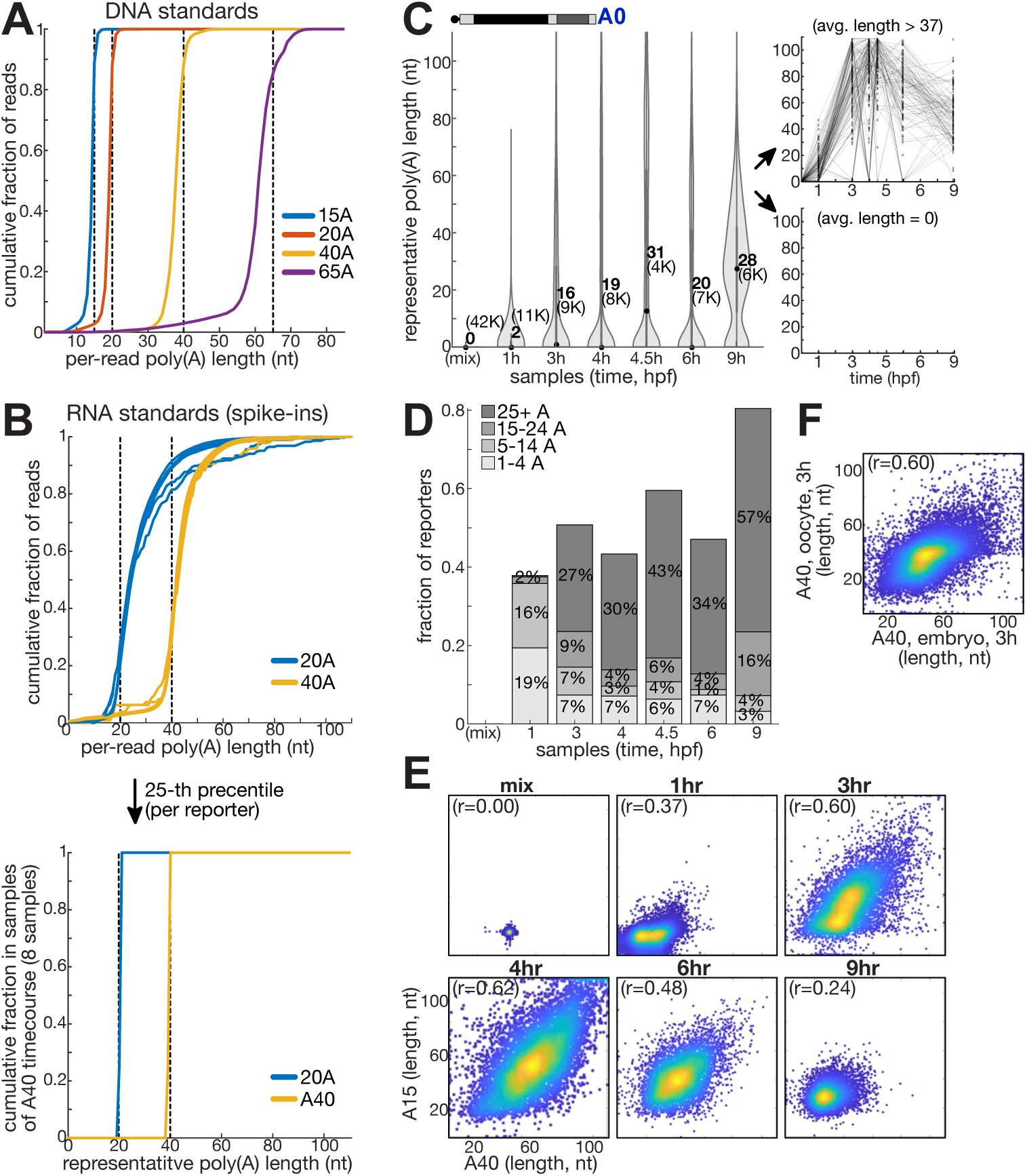
Analysis of poly(A) tail lengths by *multi-UTR*. **(A)** Distribution (y-axis, cumulative fraction of reads) of per-read estimated poly(A) tail lengths (x-axis, nt) in DNA spike-ins with different poly(A) length. **(B)** Top: distribution (y-axis, fraction of reads) of per-read estimated poly(A) tail lengths (x-axis, nt) in RNA spike-ins with different poly(A) length. Bottom: estimated representative poly(A) length for RNA spike-ins based on the 25^th^ percentile length measured per reporter. Use of the 25^th^ percentile length as representative helped to mitigate right-skewed distributions in high-throughput sequencing of long homopolymeric tracts. **(C)** Left: distribution of 0A reporters’ representative poly(A) tail lengths (y-axis, nt, 25^th^ percentile length measured per reporter) in temporal samples (x-axis) of developing zebrafish embryos. The central dot is median; gray central box bounds are 25^th^ and 75^th^ percentiles, upper and lower limits of whiskers are 1.5× interquartile ranges. Values outside of the upper and lower limits are defined as outliers. (mix) represents the in-vitro transcribed RNA, not injected into embryos. Numbers represent mean value, and number of reporters analyzed is noted in brackets. Right: two plots showing poly(A) tail lengths (y-axis, 25^th^ percentile length measured per reporter) in temporal samples (x-axis) of a subset of reporters with average poly(A) tail lengths (across all samples) at the top (top plot) or bottom (bottom plot) 2% of the distribution. **(D)** Fraction of 0A reporters (y-axis) that gain a poly(A) tail in temporal samples (x-axis) of developing zebrafish embryos. Colors represent the estimated length of the poly(A) tail (25^th^ percentile length measured per reporter), and % out of total reporters analyzed per sample is noted. **(E)** Correlation between representative poly(A) tail lengths (nt, 25^th^ percentile length measured per reporter) estimated for reporters in temporal samples from embryos injected with two different RNA libraries: A40 (x-axis) and A15 (y-axis). Colors represent density (blue = low density; yellow = high density). **(F)** Correlation between representative poly(A) tail lengths (nt, 25^th^ percentile length measured per reporter) estimated for A40 reporters in embryos (x-axis) and oocytes (y-axis) at 3 hours after injection. Colors represent density (blue = low density; yellow = high density).

**Figure S3.**
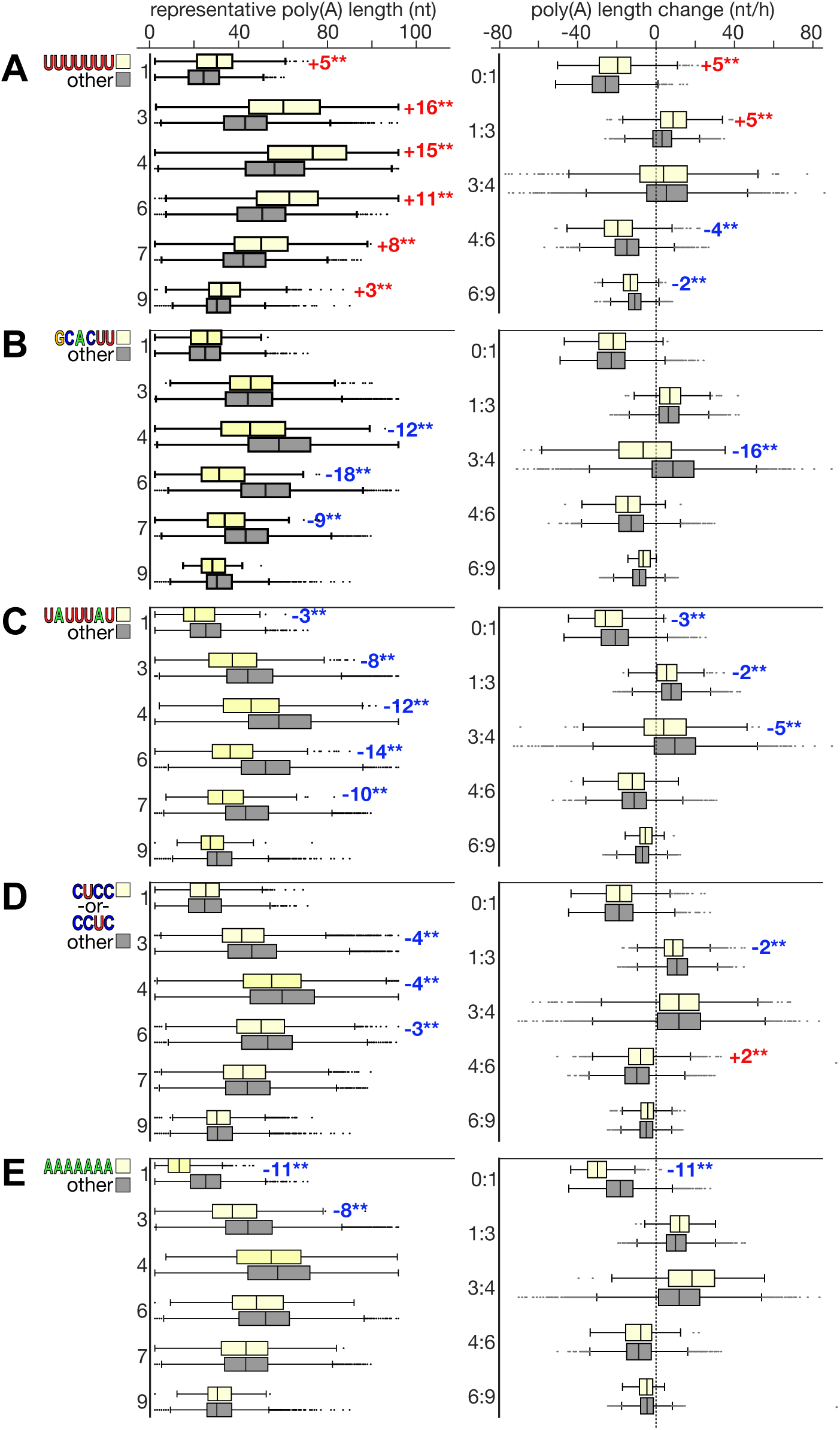
3’UTR signals associated with poly(A) length. Left: boxplots showing distribution of A40 reporters’ representative poly(A) tail lengths (x-axis, nt, 25^th^ percentile length measured per reporter) with (yellow) or without (gray) each of five specific short sequences **(A-E)**. Right: boxplots showing distribution of the temporal change in A40 reporters’ poly(A) tail lengths (x-axis, nt) with (yellow) or without (gray) a specific short sequence. Time points compared are separated by a colon (y-axis). Central line represents the median, box edges are 25^th^ and 75^th^ percentiles, whiskers extend to largest/smallest value except outliers; outlier points are plotted individually. Difference in median is noted on plots (red: higher than background, blue: lower than background), and ** denotes a significant difference by a Kolmogorov-Smirnoff test with a 1% Bonferroni multiple hypothesis correction over all tested short sequences and samples (4-7nt long, 21,760 sequences; p<10^-9^).

**Figure S4.**
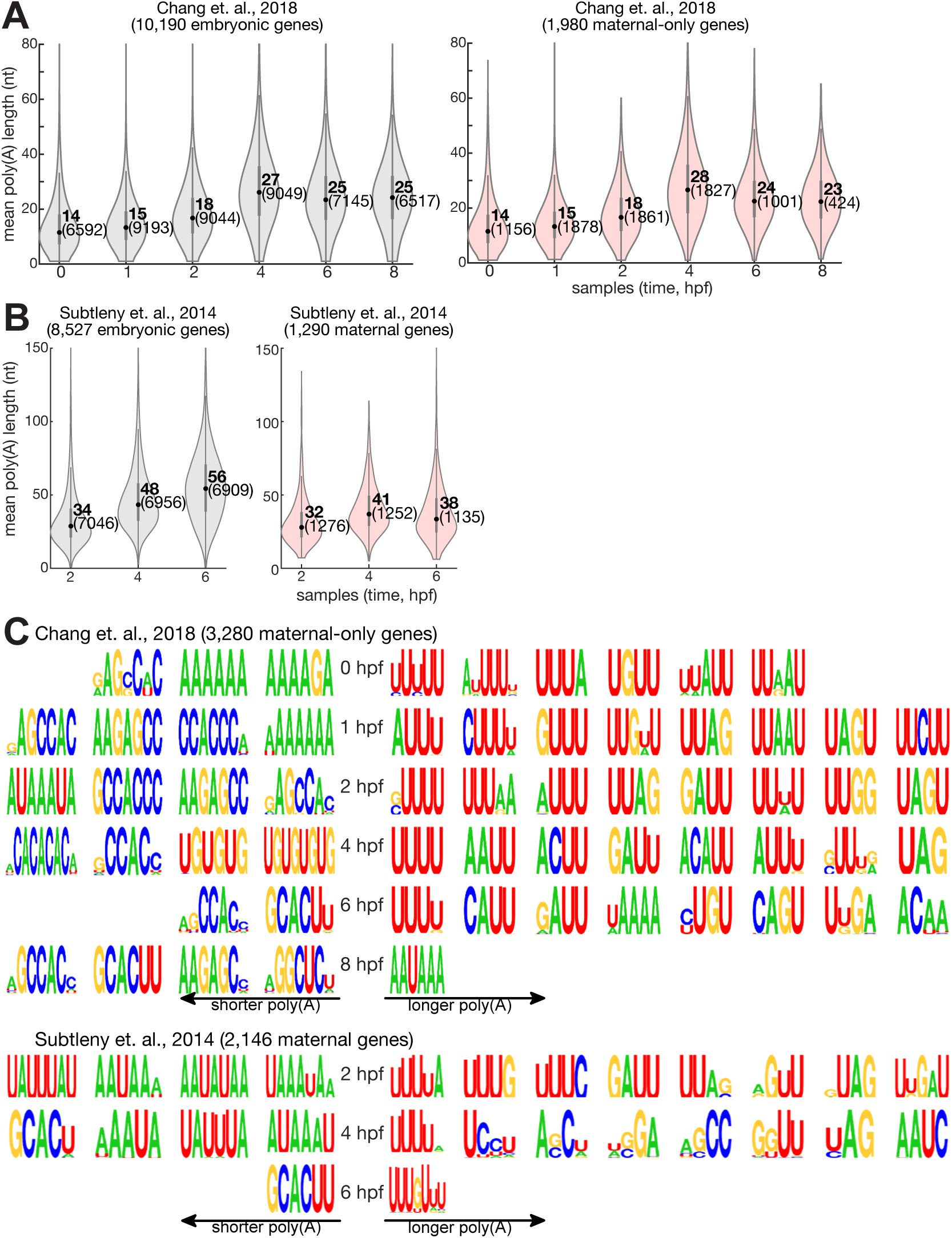
Poly(A) lengths of zebrafish native maternal mRNAs. (A-B) Distribution of native poly(A) tail lengths (y-axis, nt, mean length) in temporal samples (x-axis) of developing zebrafish embryos as measured by Chang et. al., 2018 **(A)** and Subtleny et. al. 2014 **(B)**. Left: all embryonic genes (gray). Right: only the subset of maternal genes (red). The central dot is median; gray central box bounds are 25^th^ and 75^th^ percentiles, upper and lower limits of whiskers are 1.5× interquartile ranges. Values outside of the upper and lower limits are defined as outliers. Numbers represent mean value, and number of genes analyzed is noted in brackets. **(C)** Motif logos predicted by QUANTA to represent k-mers associated with longer (right) or shorter (left) poly(A) tail lengths. Motifs that are associated with 10% or more of k-mer positions are shown.

**Figure S5.**
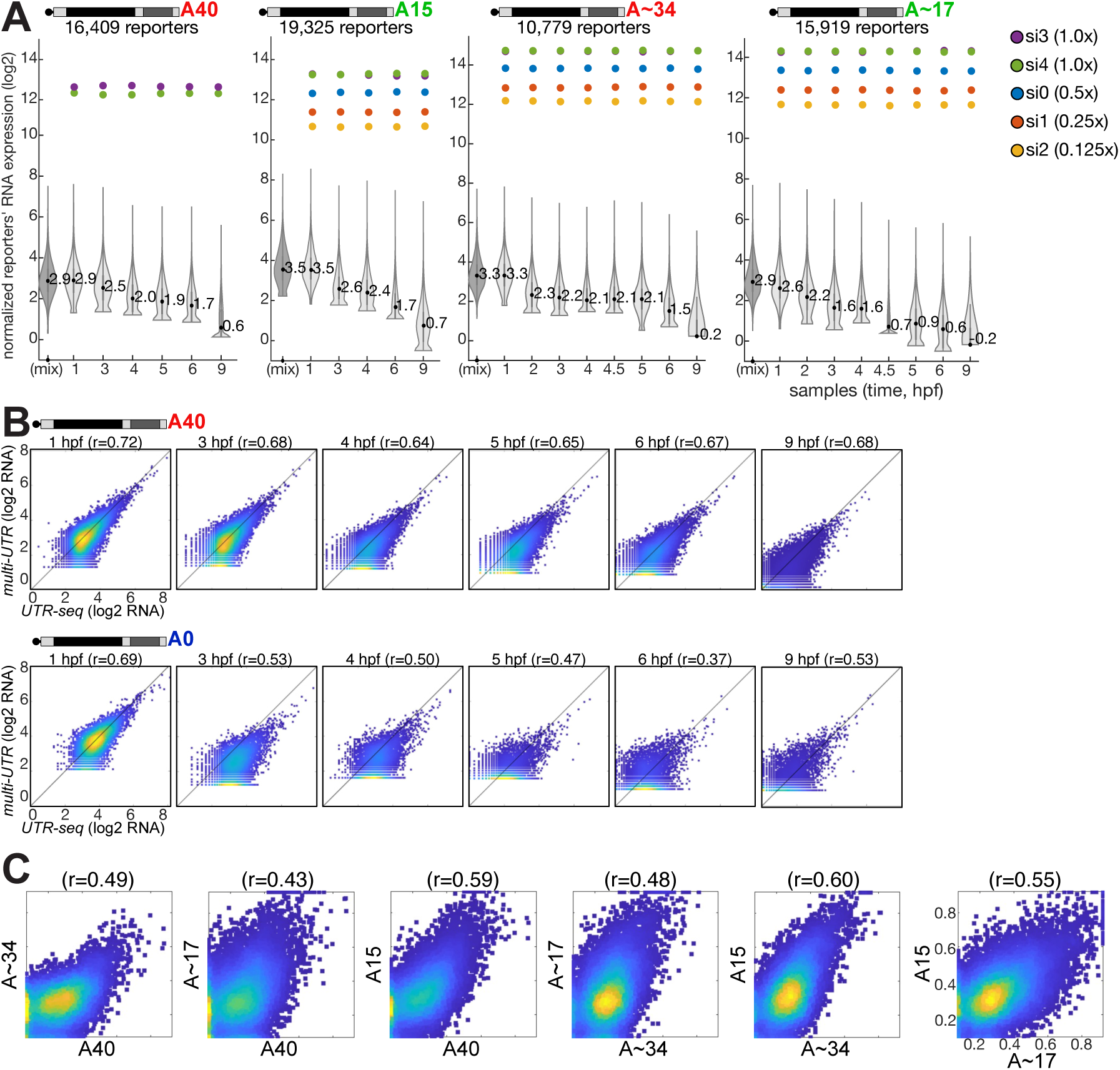
mRNA abundance quantification in *multi-UTR*. **(A)** Distribution of normalized reporter expression levels (y-axis, log2) in temporal samples (x-axis). Mean value is indicated. The central dot is median; gray central box bounds are 25^th^ and 75^th^ percentiles, upper and lower limits of whiskers are 1.5× interquartile ranges. Values outside of the upper and lower limits are defined as outliers. A set of 5 spike-ins at 2-fold incremental levels was used for normalization (colored dots). In the 40A sample, only 2 spike-ins were used for normalization. **(B)** Correlation between reporters’ expression quantification by *UTR-Seq* (x-axis) and *multi-UTR* (y-axis) in A40 reporters (top) and A0 reporters (bottom). Pearson R is indicated on top. Colors represent density (blue = low density; yellow = high density). **(C)** Correlation between degradation rates (1/h) estimated from the full temporal timecourse in different *multi-UTR* libraries (as indicated in each plot). Colors represent density (blue = low density; yellow = high density).

**Figure S6.**
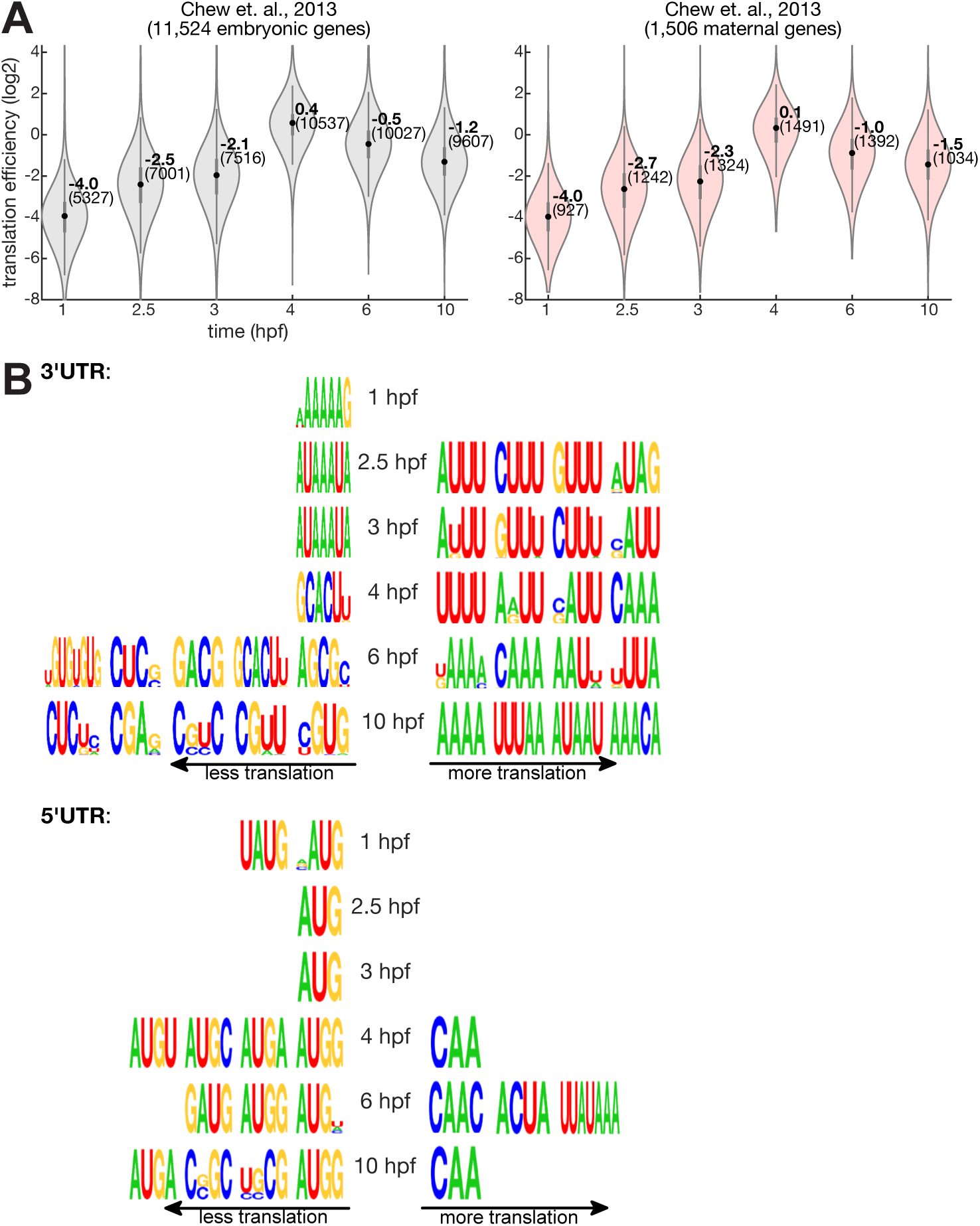
Translation efficiencies of zebrafish native maternal mRNAs. **(A)** Distribution of translation efficiencies of native embryonic genes (y-axis) in temporal samples (x-axis) of developing zebrafish embryos as measured by Chew et. al., 2013. Translation efficiency was calculated as the ratio between normalized read coverage measured in matching ribo-seq (ribosome protected fragments) and rna-seq datasets. Left: all embryonic genes (gray). Right: only the subset of maternal genes (red). The central dot is median; gray central box bounds are 25^th^ and 75^th^ percentiles, upper and lower limits of whiskers are 1.5× interquartile ranges. Values outside of the upper and lower limits are defined as outliers. Numbers represent mean value, and number of genes analyzed is noted in brackets. **(B)** Motif logos predicted by QUANTA to represent k-mers in 3’UTR (top) or 5’UTR (bottom) sequences that were associated with lower (right) or higher (left) translation efficiency. Motifs that are associated with 10% or more of k-mer positions are shown.

**Figure S7.**
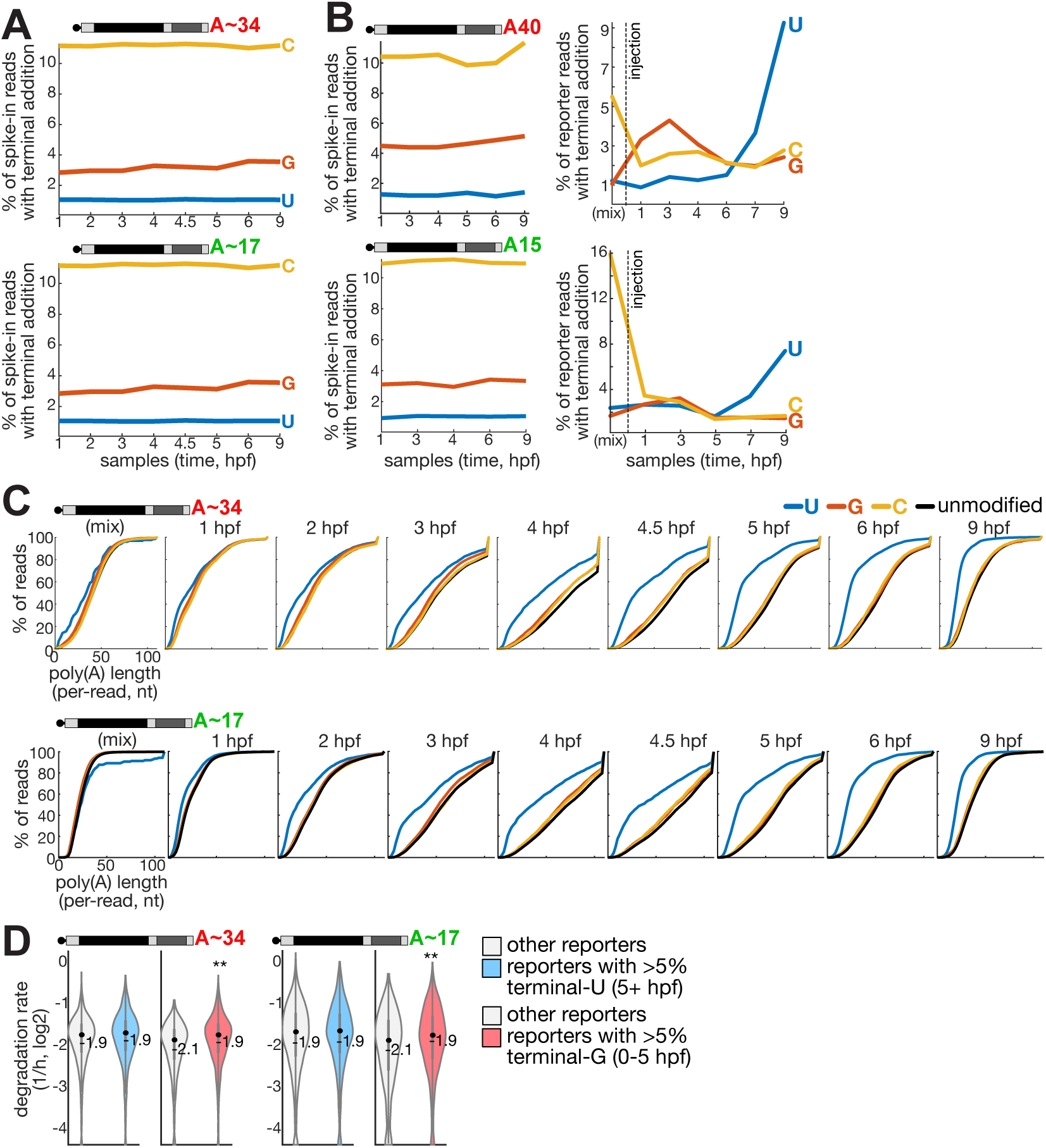
Validations of terminal modification analysis in *multi-UTR*. **(A)** Fraction of spike-in reads with terminal nucleotide additions (y-axis, 3 nucleotides: G = red, C = yellow, U = blue) in temporal samples from zebrafish embryos (x-axis) that were injected with each of two reporter libraries enzymatically polyadenylated in-vitro with poly(A) polymerase (top: A∼34, bottom: A∼17). Although library reporters were polyadenylated in-vitro by poly(A) polymerase, spike-ins were synthesized with a DNA-encoded poly(A), and therefore contain a high fraction of non-A terminal bases, serving as a control for terminal base analysis. **(B)** Fraction reads with terminal nucleotide additions (y-axis, 3 nucleotides: G = red, C = yellow, U = blue) in temporal samples from zebrafish embryos (x-axis) estimated in two reporter libraries with DNA-encoded poly(A) tails (top: A40, bottom: A15). Left: for added spike-ins, right: for reporters. **(C)** Cumulative distribution (y-axis, cumulative % of reads) of per-read poly(A) tail length (x-axis, nt) in reads with terminal nucleotide additions (G = red, C = yellow, U = blue) or without any terminal addition (black) in two ireporter libraries enzymatically polyadenylated in-vitro with poly(A) polymerase (top: A∼34, bottom: A∼17). Plots represent temporal samples, as noted. **(D)** Distribution of reporters’ degradation rates (y-axis, 1/h, log_2_), in reporters with >5% reads with terminal nucleotide additions (blue = U late (5+ hpf), red = G early (0-5 hpf)) or reporters with a lower % (gray). The central dot is median; gray central box bounds are 25^th^ and 75^th^ percentiles, upper and lower limits of whiskers are 1.5× interquartile ranges. Values outside of the upper and lower limits are defined as outliers. Numbers represent mean value, and ** denote a significant difference by Kolmogorov-Smirnoff test between modified and non-modified reporters, using a 1% Bonferroni correction.

**Figure S8.**
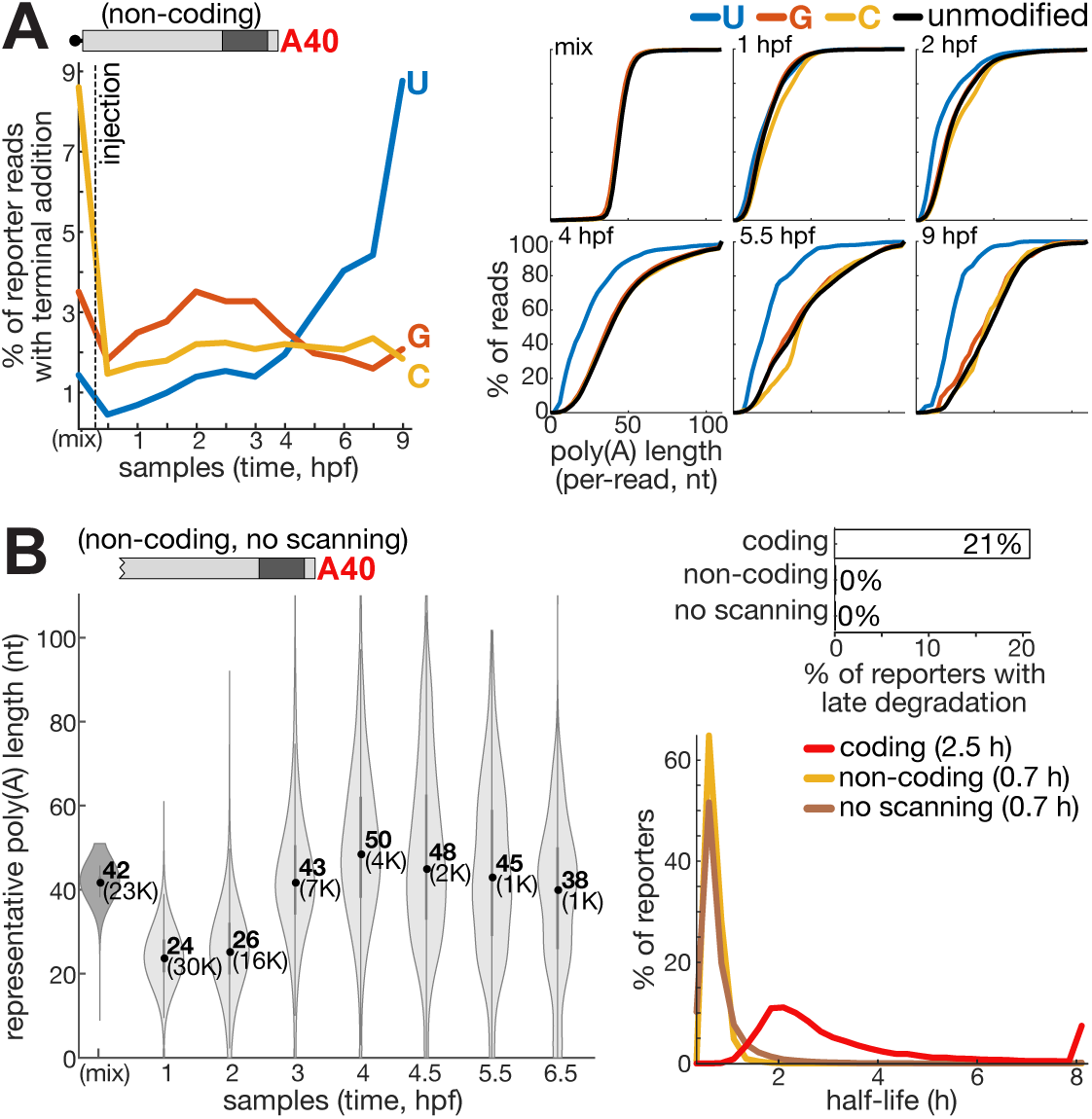
Terminal modifications of non-coding reporters. **(A)** Left: distribution of reads with terminal nucleotide additions (y-axis, 3 nucleotides: G = red, C = yellow, U = blue) in temporal samples from zebrafish embryos (x-axis) estimated in non-coding reporters. Right: cumulative distribution (y-axis, cumulative % of reads) of per-read poly(A) tail lengths (x-axis, nt) in reads with 3 terminal nucleotide additions (G = red, C = yellow, U = blue) or without any terminal addition (black) in non-coding reporters. Plots represent temporal samples, as noted. **(B)** Left: distribution of representative poly(A) tail lengths (y-axis, nt, 25^th^ percentile length measured per reporter) of non-coding reporters with a non-functional cap (ApppG), in temporal samples (x-axis) of developing zebrafish embryos. The central dot is median; gray central box bounds are 25^th^ and 75^th^ percentiles, upper and lower limits of whiskers are 1.5× interquartile ranges. Values outside of the upper and lower limits are defined as outliers. (mix): in-vitro transcribed RNA library, not injected into embryos. Numbers represent mean value, and number of reporters analyzed is noted in brackets. Right: distribution (y-axis, % of reporters) of half-lives (x-axis, h) estimated for coding and non-coding reporters with either a functional cap (“non-coding”) or with a non-functional ApppG cap (“no scanning”). Type of library and mean half-life estimated (excluding upper and lower bound values) is noted in legend. Top histogram shows % of reporters in each library with late-onset of degradation, identified based on early stability (half-life ≥10 h, estimated on early samples (1-4 hpf)) and late degradation (half-life <10 h, estimated on late samples (4-9 hpf)).

**Figure S9.**
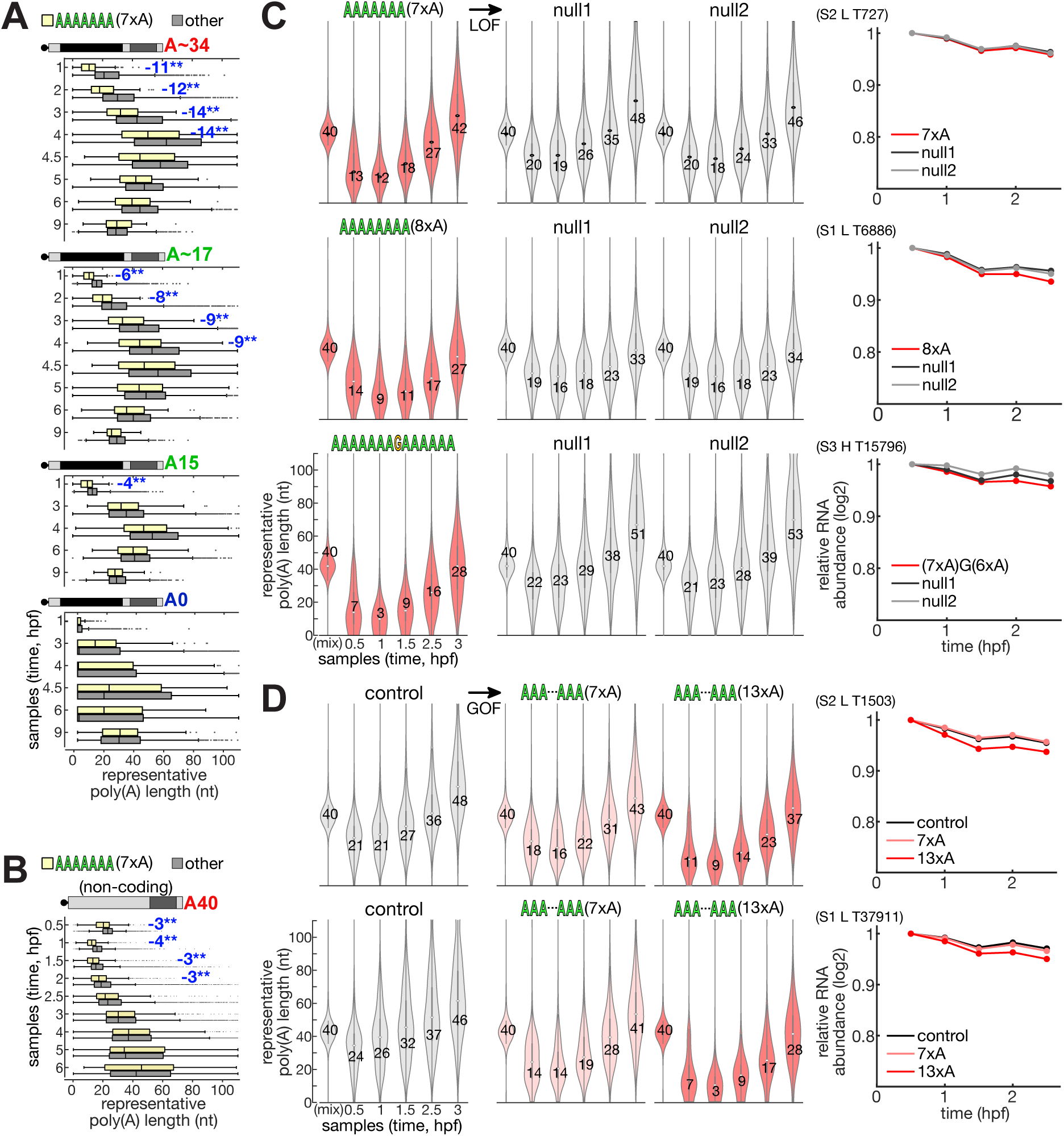
Further analysis and validation of the internal 3’UTR A-tract motif. (A-B) Distribution of representative poly(A) tail lengths (x-axis, nt, 25^th^ percentile length measured per reporter) with (yellow) or without (gray) an A-tract motif. Central line represents the median, box edges are 25^th^ and 75^th^ percentiles, whiskers extend to largest/smallest value except outliers; outlier points are plotted individually. Difference in median is noted on plot (blue = lower than background), and ** denotes a significant difference by a Kolmogorov-Smirnoff test with a 1% Bonferroni multiple hypothesis correction over all tested short sequences and samples (4-7nt long, 21,760 sequences; p<10^-9^). **(A)** For coding reporter libraries with different initial poly(A) lengths. **(B)** For a non-coding reporter library. **(C-D)** Left: distribution of representative poly(A) tail lengths (y-axis, nt, 25^th^ percentile length measured per reporter) of validation reporters with or without specific A-tract motifs, as noted on top of each plot, in temporal samples (x-axis) of developing zebrafish embryos. The central dot is median; gray central box bounds are 25^th^ and 75^th^ percentiles, upper and lower limits of whiskers are 1.5× interquartile ranges. Values outside of the upper and lower limits are defined as outliers. Numbers represent mean values. Gray: validation reporters not containing an A-tract motif; light red: weak A-tract motifs; dark red: strong A-tract motifs. Right: temporal (x-axis, time, hpf) RNA abundance relative to the first timepoint (y-axis) of the reporters shown on the left panel (colors as annotated in legend). **(C)** Loss-of-function (LOF) perturbations of 3 reporters with a strong A-tract motif that is mutated in two different ways (null1 and null2). **(D)** Gain-of-function (GOF) mutations creating A-tract motifs of medium (7xA) and long (13xA) length in two background control reporters that did not contain the motif originally.

Supplementary Table S1: Full list of k-mers associated with poly(A) tail lengths and grouped into 9 clusters.

Supplementary Table S2: Full list of k-mers associated with degradation rates and grouped into 9 clusters.

Supplementary Table S3: Full list of k-mers associated with terminal modifications and grouped into 9 clusters.

Supplementary Table S4: List of primer and other sequences used in this study.

## Notes

### Competing Interest Statement

The authors have declared no competing interest.

