## Supplemental Table 4 for "3’UTR-directed control of poly(A) tail dynamics and mRNA stability in vertebrate embryos"

### Table S4: Oligonucleotides used in the study

#### Primers for generation of transcription templates

**216.SP6UTRseqF**
5'-ATTTAGGTGACACTATAGAATACAC-3' 25-mer
Forward primer for amplification of template for SP6 transcription of UTR-seq reporters.

**222.SP6UTRseqFdU**
5'-ATTTAGGTGACAC/ideoxyU/ATAGAATACAC-3' 25-mer
Forward primer for generation of templates for SP6 transcription of UTR-seq reporters at high salt conditions. Contains deoxyuridine, which can be removed with USER enzyme after PCR.

**223.UTRseqSpeI**
5'-ACTAGTGTAATACGACTCACTATAG–3' 25-mer
Reverse primer for generation of templates for SP6 transcription of 0(A) UTR-seq reporters. Includes complete SpeI site.

**273.15A_SpeI**
5'-TTTTTTTTTTTTTTTACTAGTGTAATACGACTC-3' 33-mer
For construction of UTR-seq transcription template with 15(A).

**300.DS_40A**
5'-TTTTTTTTTTTTTTTTTTTTTTTTTTTTTTTTTTTTTTTTACTAGTGTAATACGACTC-3' 58-mer
Reverse primer for generation of UTR-seq transcription template with 40(A) by PCR.

#### RNA 3’ end capture oligos

**272.15A_capture**
5'-TTTTTTTTTTTTTTTACTAGTGTAA/3AmMO/-3' 25-mer
For sequestering of RNA 3' end in UTR-seq with 15A

**350.0A_capture6th**
5'-ACTAGTGTAATACGACT/3AmMO/-3' 17-mer
For sequestering of RNA 3' end in UTR-seq with 0(A), ending with **6th** nucleotide of SpeI site.

**Library amplification primers**

**Downstream_P5b_F_C2_R**
5'-AATGATACGGCGACCACCGAGATCTACACTCTTTCCCTACACGACGCTCTTCC-3’
UTR-seq downstream primer. Read1 reads from RT side. Does not carry an index.

**Indexed downstream PCR primer**
5’-AATGATACGGCGACCACCGAGATCTACACNNNNNNNNACACTCTTTCCCTACACGACGCTCTTCC-3’
P5
NNNNNNNN- i5 (index 2)
Partial Read1

**Upstream library amplification primer (indexed UTR-seq primers from plate)**
5’-CAAGCAGAAGACGGCATACGAGATNNNNNNNGTGACTGGAGTTCAGACGTGTGCTCTTCCGATCTGGAGATCTGAGTTCAAGGAT-3’
P7 reverse complement
NNNNNNN- i7 (index 1)
Read2 reverse complement
const_1

**Primers for generating DNA standards**

**274.standards.R**
5'-AATGATACGGCGACCACCGAGATCTACACTCTTTCCCTACACGACGCTCTTCCGATCTCTGCGGCCGCTCTTC-3' 73-mer
Reverse PCR primer for generation of DNA standards for poly(A) sequencing from spike-in plasmids. Modified from UTR-seq downstream primer.
P5
Full Read1 primer
Anneals to sequence downstream of poly(A)in spike-in plasmids

| |
|---|
| **275.15A.stand.R**<br>5'-CTGCGGCCGCTCTTCTTTTTTTTTTTTTTTACTAGTGTAATACGAC-3' 46-mer<br>Reverse PCR #1 primer for generation of a 15(A) DNA standard for poly(A) sequencing. Use on plasmid with long poly(A) to reduce and allow detection of annealing of PCR handle (green) to plasmid.<br>PCR handle for primer #274<br>Anneals to sequence upstream of poly(A) and to poly(A) |
| **Primers for generation of mutated reporters**<br><br>**397.S3_H_T15796.F**<br>5'-ggagatctgagttcaaggatATCTACCGTGTGTAAGAGATATTCTATAATCTATATAGTTTCATTCCATATTG-3'<br>Forward primer for modification of polyA motif in reporter S3_H_T15796<br><br>**398.S3_H_T15796.M1**<br>5'-GATTATACAGTATAGAAATGTGCGATACTACTGAAACTCATAAACAATATGGAATGAAACTATATAGATTATAG-3'<br>Middle primer for generation of mutation M1 of polyA motif in reporter S3_H_T15796<br><br>**399.S3_H_T15796.M2**<br>5'-GATTATACAGTATAGAAATGTGCGATAACTAGTACTCGACTAAACAATATGGAATGAAACTATATAGATTATAG-3'<br>Middle primer for generation of mutation M2 of polyA motif in reporter S3_H_T15796<br><br>**400.S3_H_T15796.R**<br>5'-gactcactatagttctagatCAATGCCCAAACAGATTATACAGTATAGAAATGTGCG-3'<br>Reverse primer for modification of polyA motif in reporter S3_H_T15796<br><br>**401.S2_L_T727.F_M1**<br>5'-ggagatctgagttcaaggatCAGTGCTTATTGAGATTGAAATAAGATT**TCAGTAG**TTGAAATTGAGATGAGAAATATCGT-3'<br>Forward primer for modification of polyA motif in reporter S2_L_T727 (mutation M1)<br><br>**402.S2_L_T727.F_M2**<br>5'-ggagatctgagttcaaggatCAGTGCTTATTGAGATTGAAATAAGATT**TACTAGT**TTGAAATTGAGATGAGAAATATCGT-3'<br>Forward primer for modification of polyA motif in reporter S2_L_T727 (mutation M2) |

**403.S2_L_T727.Mid**
5'-CAACAATCGTTATTTCCTAAAGCAGTCACAAGTGTTGATATAACGATATTTCTCATCTCAATTTC-3'
Middle primer for modification of polyA motif in reporter S2_L_T727

**404.S2_L_T727.R**
5'-gactcactatagttctagatGACGTCCACAACAATCGTTATTTCCTAAAG-3'
Reverse primer for modification of polyA motif in reporter S2_L_T727

**405.S1_L_T6886.F**
5'-ggagatctgagttcaaggatAACTGAGGACCTGTGCTAAATCAATTGGTCTGGACTCTCAGAATTGTCTCATTACAGATG-3'
Forward primer for modification of polyA motif in reporter S1_L_T6886

**406.S1_L_T6886.M1**
5'-GAGAAACGACACTTTGAGCTTTGGGGTACAAGGCA**CTACTGAA**ACATCTGTAATGAGACAATTCTG-3'
Middle primer for generation of mutation M1 of polyA motif in reporter S1_L_T6886

**407.S1_L_T6886.M2**
5'-GAGAAACGACACTTTGAGCTTTGGGGTACAAGGCA**ACTAGTAC**ACATCTGTAATGAGACAATTCTG-3'
Middle primer for generation of mutation M2 of polyA motif in reporter S1_L_T6886

**408.S1_L_T6886.R**
5'-gactcactatagttctagatGGAAATGAGAAACGACACTTTGAGC-3'
Reverse primer for modification of polyA motif in reporter S1_L_T6886

**409.S1_L_T37911.F**
5'-ggagatctgagttcaaggatACATCTTCACTCCATGTTGTTTATTCTCTGGAATGTATTCATTTTAAATTGTAG-3'
Forward primer for adding a polyA motif to reporter S1_L_T37911

**410.S1_L_T37911.7A**
5'-GAATATTTGCTTGAAATAATTCAC**TTTTTTT**AGCATTAAGATAAATTAAAAACTACAATTTAAAATGAATACATTCCAG-3'

Middle primer for adding a polyA motif to reporter S1_L_T37911 (7A)

**411.S1_L_T37911.13**
5’-GAATATTTGCTTGAAATAATTCAC**TTTTTTTTTTTTT**AAGATAAATTAAAAACTACAATTTAAAATGAATACATTCCAG-3’
Middle primer for adding a polyA motif to reporter S1_L_T37911 (13A)

**412.S1_L_T37911.R**
5’-gactcactatagttctagatTGCAGAATATTTGCTTGAAATAATTCAC-3’
Reverse primer for adding a polyA motif to reporter S1_L_T37911

**413.S2_L_T1503.F**
5’-ggagatctgagttcaaggatTTGATCACAATTCCCTTGTTTCTTGTTTCTAGTAGTTCTTACTAGTTCTCACAGATAG-3’
Forward primer for adding a polyA motif to reporter S2_L_T1503

**414.S2_L_T1503.7A**
5’-CTGGTAATTAATCATTATGATAATGATGTTTTTTTAATTCAGATACTATCTGTGAGAACTAGTAAGAAC-3’
Middle primer for adding a polyA motif to reporter S2_L_T1503 (7A)

**415.S2_L_T1503.13A**
5’-CTGGTAATTAATCATTATGATAATGATGTTTTTTTTTTTTTGATACTATCTGTGAGAACTAGTAAGAAC-3’
Middle primer for adding a polyA motif to reporter S2_L_T1503 (13A)

**416.S2_L_T1503.R**
5’-gactcactatagttctagatAATAGCCCTGGTAATTAATCATTATGATAATGATG-3’
Reverse primer for adding a polyA motif to reporter S2_L_T1503 (13A)

**419.S3_H_T15796.wt**
5’-GATTATACAGTATAGAAATGTGCGATATTTTTTCTTTTTTTAAACAATATGGAATGAAACTATATAGATTATAG-3’ 74-mer
Middle primer for generation of wild type UTR-seq reporter S3_H_T15796.wt

**420.S2_L_T727.F_wt**

5’-ggagatctgagttcaaggatCAGTGCTTATTGAGATTGAAATAAGATT**AAAAAAA**TTGAAATTGAGATGAGAAATATCGT-3’ 80-mer
Forward primer for generation of wild type UTR-seq reporter S2_L_T727.F

**421.S1_L_T6886.wt**
5’-GAGAAACGACACTTTGAGCTTTGGGGTACAAGGCA**TTTTTTTT**ACATCTGTAATGAGACAATTCTG-3’ 66-mer
Middle primer for generation of wild type UTR-seq reporter S1_L_T6886

**422.S1_L_T37911.wt**
5’-GAATATTTGCTTGAAATAATTCAC**TTTAATAAGCATT**AAGATAAATTAAAAACTACAATTTAAAATGAATACATTCCAG-3’ 79-mer
Middle primer for generation of wild type UTR-seq reporter S1_L_T37911

**423.S2_L_T1503.wt**
5’-CTGGTAATTAATCATTATGATAATGATG**TAATTCTAATTCA**GATACTATCTGTGAGAACTAGTAAGAAC-3' 69-mer
Middle primer for generation of wild type UTR-seq reporter S2_L_T1503

**Primers for adding 15 bases of flanking sequence to cloned reporters, for Gibson assembly**

**sfGFP_const1_F+15**
5’-AGCTCGAGGATGCTAGGAGATCTGAGTTCAAGGAT-3'

**sfGFP_const2_R+15**
5’-TTACTAGTGTAATACGACTCACTATAGTTCTAGAT-3’

**296.RNA3pAdap_rAPP**
5'–/5rApp/CGCGAAGTNNNNNNNNNNNNAGGTAGATCGGAAGAGC/3AmMO/–3' 37-mer
Adapter to be ligated to the 3' end of RNAs for reverse transcription.

**239.RNAadapterRT1**
5'-ctacacgacgctcttccgatctGCTCTTCCGATCTACCT-3' 39-mer
RT primer for use with 296.RNA3pAdap_rAPP.
lowercase letters represent 3' end of Read1 primer (same as UTR-seq RT primer handle). Underlined 17 nt are target sequence of UTR-seq downstream primers.

UPPERCASE LETTERS represent sequence that anneals to 3' side of 296.RNA3pAdap_rAPP.

**UTR-seq RT primer**

5’-**CTACACGACGCTCTTCC**GATCTNNNNNNNNGACTCACTATAGTTCTAGAT-3’

Partial Read1. Target sequence of UTR-seq downstream primers is in bold.

NNNNNNNN- UMI

Reverse-complement of const_2

**3’UTR sequences of added spike ins (including upstream and downstream constant regions, labeled in blue)**

**Spike in 0**

**GGAGATCTGAGTTCAAGGAT**TTGCTTAGAAGCGCGGTGGAACACCATGTAGTGCTTCCCTGTTATCTAGCTATATTAGGTCCAAGGTTTCTAAGTTGTAA
CAATTAGTGCCGTAACTACCACAGCAGGTG**ATCTAGAACTATAGTGAGTC**

**Spike in 1**

**GGAGATCTGAGTTCAAGGAT**CATAACGCACGCAGCAAGAACTATCATTTGATTCGACGCCGCTTAAACCGATCTTAGTTATGGGCCATCGTAAGTAATCG
GGCTTGTTTCCGCGTGTATCGTCTGAAGTT**ATCTAGAACTATAGTGAGTC**

**Spike in 2**

**GGAGATCTGAGTTCAAGGAT**ACCAGCGGCTTCGGATAGATGGGAGATATCGATTGTCTGAGAGCTGTGCTATTCCTTTTATAAACGTAGTTAACGCTATA
TCCGCAAGTTAAAGGCTACGCACGCATTTA**ATCTAGAACTATAGTGAGTC**

**Spike in 3**

**GGAGATCTGAGTTCAAGGAT**AATAGGGTAGGCATTCTACGCTCCGTTCTATGATCGTCGGCCACATATTAAAGCATTTTTCTGACAACTTTAACAAAACG
TGTAACTTGCACGGCCGAACGCTCATTGCT**ATCTAGAACTATAGTGAGTC**

**Spike in 4**

**GGAGATCTGAGTTCAAGGAT**AGGGGGCCCTGTAGGGACTGTCATGAAGATTATTAGTCGTGAAGATTCATGGGGAATTTGTCTGTCGAATCGAAATGTTT
ACCCTGCTTATCGAGGCCGTGATCGCGCAT**ATCTAGAACTATAGTGAGTC**
